# Imaging of specialized plant cell walls by improved cryo-CLEM and cryo-electron tomography

**DOI:** 10.1101/2025.06.01.657137

**Authors:** J Daraspe, E Bellani, D De Bellis, C Genoud, N Geldner

## Abstract

Cryo-focused ion beam scanning electron microscopy (cryo-FIBSEM) has become essential for preparing electron-transparent lamellae from cryo-plunged and high-pressure frozen specimens. However, targeting specific cellular features within large, complex organs remains challenging. Here we present a series of technical improvements significantly enhancing the efficiency and accessibility of the Serial Lift-Out and SOLIST (Serialized On-grid Lift-In Sectioning for Tomography) procedures that are revolutionizing the field. We were able to extend the cryo-FIBSEM session from 24 hours to 5 days without interruptions. In addition, we describe a modified silver-plated EasyLift^TM^ needle that eliminates the need of the copper or gold block between the original tungsten needle and the sample. Moreover, we describe a strategy that significantly reduces curtaining effects. Finally, we report a precise routine to target a lamella with a precision of approximately 1 µm in X,Y and Z. Together, these modifications considerably reduce contamination risk and preparation time, making cryo-lift-out techniques more accessible for routine structural biology applications on any type of tissue. Here, we demonstrate the power of our technique by targeting several specific wall structures that are of crucial importance for root function in plants and that were previously inaccessible to cryo-electron tomography (cryo-ET). High-pressure freezing (HPF) of plant tissues presents unique challenges for cryo-electron microscopy sample preparation due to the overall sample size, the individual cells size, their rigid cell wall and finally, their large vacuoles, which contain large amounts of rather diluted water solutions compared to cytosol. The internal root structures targeted are the Casparian strip (CS), suberin lamellae (SL), as well as secondary wall of xylem vessels, requiring reaching a targeting precision of 5 μm in a 3 mm long and 80-120 μm thick root tip. Our technological improvements for the cryo-correlative light and electron microscopy (cryo-CLEM) workflow enabled successful, targeted cryo-ET in plant roots. We noticed that, despite ice formation in vacuoles and to some degree in the cytosol, the plasma membranes and cell walls are remarkably well preserved, providing stunning insights into the native, hydrated nano-structure of plant cell walls, previously only observable with contrasting agents and in a dehydrated state.

## Introduction

Cryo-electron microscopy (cryo-EM) has revolutionized structural biology, allowing to study molecular complexes at near-atomic resolution ^1^. There is now a shift in focus toward cryo-electron tomography (cryo-ET)^2^, as it allows to image cell ultrastructure in their most native state. At the same time, it can enable imaging of macromolecules within intact cells or tissues by capturing a series of tilted projections and reconstructing three-dimensional volumes^2^ ^3^.

Molecules can be directly imaged on grids in the cryo-transmission electron microscope. Larger samples, by contrast, stop electrons and cannot be imaged without being thinned^3^. Preparing thin, electron-transparent sections from complex samples remains challenging. Cryo-ultramicrotomy^4^, as developed by Dubochet et al., exists for 40 years, but is difficult to apply in tissue ^5^. Furthermore, targeting small and cell-type specific regions, such as Casparian strips in plant roots is so difficult and prone to chance that it has never been done. Cryo-FIBSEM has been developed to cut and thin cryo-lamellae on the grid in which the sample is frozen. This first approach has allowed to directly thin out adherent cells, bacteria and yeasts successfully^6^. Yet, plunge-freezing is limited to thin samples below a couple of micrometers. Any more complex tissues, organoids or large cells cannot be frozen and thinned out directly on the grid. The only available method to vitrify larger samples is the HPF. It requires placing a small piece of tissue (less than 200 μm thick) inside a carrier and insert it into a vitrifying instrument. In such cases, the final product is a piece of vitrified tissue inside a carrier, meaning it is not suitable to be used directly in cryo-TEM. The waffle method^7^ consists of placing the sample on a grid inside the carrier before freezing. The grid is then collected and placed into a cryo-FIBSEM for thinning to obtain cryo-lamellae. While this approach is effective for specimens smaller than the grid bar thickness (typically 20-25 µm) and can accommodate slightly larger samples with the addition of a spacer, it presents significant limitations for larger organisms or tissue samples. It is not suitable for our work with extensive root systems, as the dimensional constraints of the grid would introduce sample damage. Recent advances such as Serial Lift-Out^8^ and SOLIST^9^ have improved throughput through innovative milling and lamella attachment strategies^10^. This workflow allows to high-pressure freeze pieces of tissue in a carrier and then transfer a region of interest on a grid to carve cryo-lamellae suitable for cryo-ET. This approach allows preparation of thin sections from virtually any part of a sample, including dense tissues and complex cellular structures.

Yet, the underlying technical constraints of cooling system stability and sample manipulation continue to limit routine adoption of these methods. Despite having successfully implemented these above workflows, we had to overcome major hurdles to apply these methods to target specific cell wall structures in plant roots.

While cryo-lift-out methods have emerged as powerful solutions for accessing these samples^10^, their practical implementation has been constrained by technical limitations. Our improved protocols and workflow allow us to considerably extend the time of the cryo-FIBSEM session, avoid the use of intermediary copper block on the needle, improve the targeting precision between the cryo-fluorescence microscope and the cryo-FIBSEM and reduce curtaining effect on final lamellae. Here, we demonstrate how these improvements allowed us to target CS in *Arabidopsis thaliana* roots. CS are a network of 2-3 μm thin cell wall impregnations, analogous to animal tight junctions, and which are formed in a single cell layer at a depth of 50 μm in the 80-120 μm diameter root and at approximately 2 mm from the root tip.

Our significant workflow improvements render these powerful techniques much more accessible and are significantly broadening the applicability of the techniques to solve new biological questions that were out of reach.

## Results

### Meristematic cells

Tomograms from cryo-lift-out of root meristematic cells generally showed the best structural conservation with the least degree of ice crystals formation^11^ (Fig. 1A-C, movie 1-2). This can be explained by the high cytoplasmic density, absence of large central vacuole and thinner, less rigid walls of these undifferentiated actively dividing cell populations in the root tip. Fig. 1B (movie 1) shows an example of a meristematic cell with a well-preserved Golgi stack, a multi-vesicular body and a mitochondrion as well as large numbers of ribosomes. Even in meristematic cells, chemical fixation in classical TEM protocols induces protoplast shrinkage, leading to plasma membrane detachment and invaginations^11^. Due to these artifacts, regularly observed extracellular membrane structures in classical TEM^11^ appeared as very heterogenous structures that were often considered artifacts. It is therefore interesting to observe smoothly shaped extracellular membranes in our cryopreserved samples (Fig. 1C, movie 2). The large extracellular vesicle or tubule is observed at a corner between three cells, associated with an invagination of the plasma membrane, which clearly surrounds the extracellular membrane. The origin of such structures in plants remains unclear, but they should arise from intracellular double-membrane structures, such as autophagosomes, multi-vesicular bodies, or various other less-known compartments such as EXPO^12,13^. Interestingly, the stamp-like shape of the extracellular membrane suggests that it becomes appressed against the wall, possibly due to the pressure exerted by the protoplast. Such flattened structures are also something we observe in extracellular membranes during CS formation (see below). Other features repeatedly observed in TEM images are electron-dense, precipitation-like structures within the plant cell wall. Such structures were difficult to interpret, since precipitates of this type can easily arise from heavy metals used for staining in room temperature TEM. We were therefore intrigued to repeatedly observe such precipitates in our cryo-fixed lamellae in the absence of any added heavy metals (Fig. 1C, movie 2). Due to their enhanced presence in cell corners and in positions near the middle lamella (see below), we speculate that these structures might arise from the presence of the high-levels of calcium in the pectin-rich regions of the cell wall.

**Figure 1.**
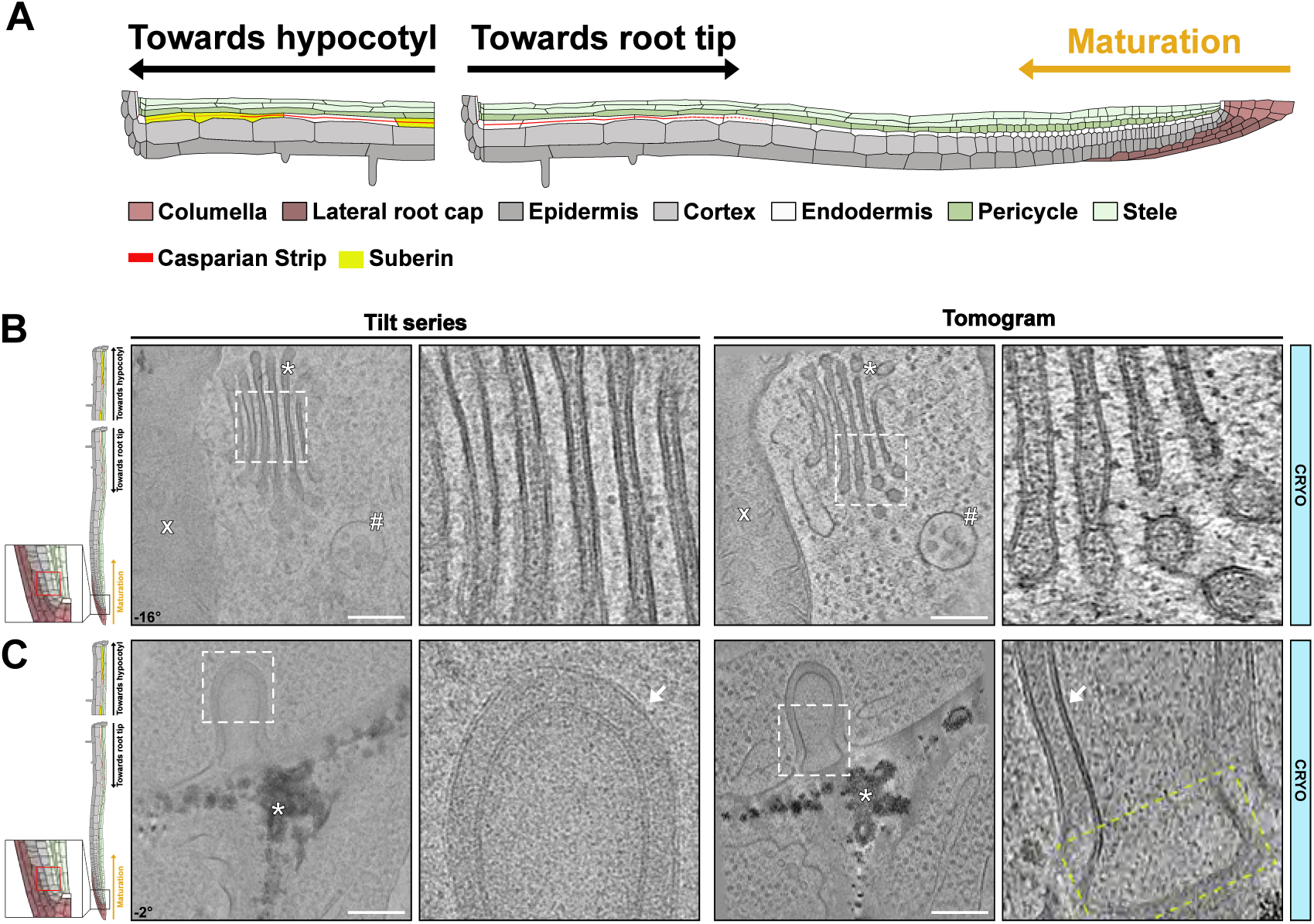
Root tip cryopreservation and organelles identification. **A)** Overview of *Arabidopsis thaliana* root, indicating differentiation gradients and cell types. Localization from which images have been taken are shown based on this schematic for all figures. **B**) Example of images obtained from the root tip. White asterisk - Golgi apparatus. White x - Mitochondria. White # - Multivesicular body. Movie 1 **C**) Example of images obtained from the root tip. Presence of a large extracelluar vesicle with invagination of plasma membrane. White asterisk - Electron-dense precipitates (Movie 2). White arrow – well preserved plasma membrane. Yellow rectangle – highlights stamp-like shape of the extracellular membrane. Corresponding angle from the tilt-series is shown on images. Scale bar = 200nm. Zoom-in field of view of 250nm.

### Unmodified cell walls in differentiated cells

For clarity, we will first describe unmodified cell walls of differentiated cells, before focusing on specific cell wall modifications. Despite increased ice crystals formation in cytosol and vacuoles of differentiated cells, we could observe well conserved rough ER cisternae, multi-vesicular bodies and other compartments (Fig. 2A, upper panel, movie 3). Below the plasma membrane at the cell corner, we could see well-conserved cortical arrays of microtubules (Fig. 2A, upper panel, movie 3). The interface between plasma membrane (PM) and cell wall showed surprisingly good conservation in our tomograms. Notably, we observed aligned, parallel fibrils in the wall proximal to the plasma membrane, probably representing highly ordered arrays of cellulose microfibrils. In chemically fixed, osmium stained, and resin embedded samples imaged at room temperature TEM, we could never observe the same degree of details or ordered arrangement of microfibrils. At best, walls in TEM pictures display a disorganized, vaguely fibrous structure, invariably appearing to have shrunk during dehydration. For adequate comparison, we have generated a tilt-series of resin embedded sections at the same resolution as the cryo-ET (Fig. 2A, bottom panel). In the distal parts of the wall (towards middle of the wall, more distant from the PM), the cell corner showed more disorganized microfibrils of lower density, consistent with the known, pectin-rich nature of this wall region. As observed in the meristematic cells, electron-dense precipitates were associated with cell corners and rarely observed in cell wall areas with dense and organized microfibril arrays. This further supports the idea that these precipitates are derived from pectin-associated calcium. Fig. 2A also shows the endodermal cells having a homogenous, electron-lucent layer between the fibrillar primary cell wall and the PM. This is not observed in the cortical cells. Such differences in spacing between PM and the primary wall could not have been interpreted in room temperature TEM, as it resembles the frequently occurring artefact of PM detachment induced by fixation^11^. We now suspect that such layers are associated with cell walls that are being actively synthesized, remodeled or modified.

**Figure 2.**
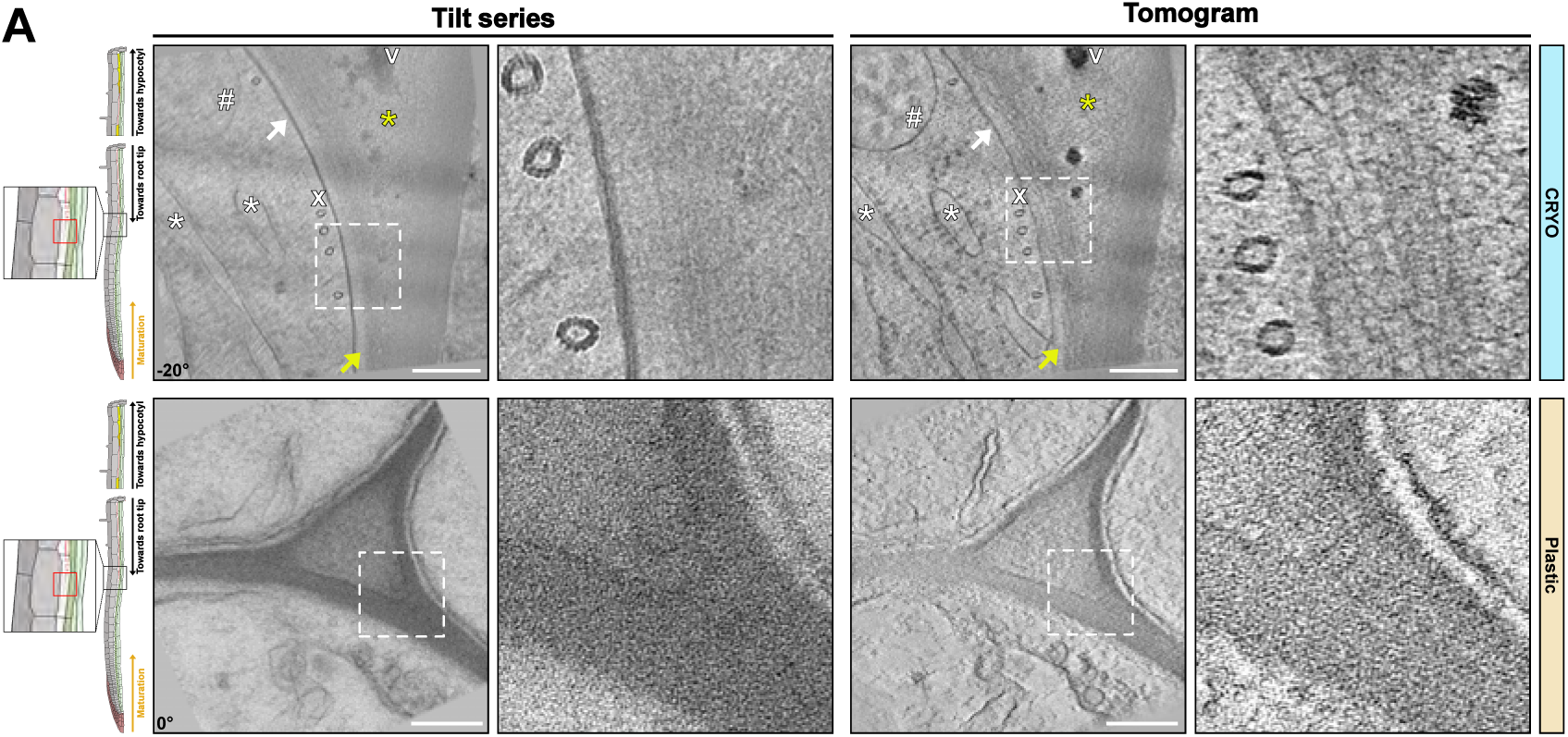
Cell wall differentiation in the early Casparian strip (CS) formation zone at approximately 1 mm from the root tip. Upper panels show the cell wall preservation in cryo-conditions (Movie 3) while the lower panels show the same area in chemically fixed, resin embedded samples. White asterisk - ER cisternae. Yellow asterisk - pectin-rich region composed of disorganized fibrils of lower density. White # - multivesicular body. White V - Electron-dense precipitates. White x - arrays of microtubules. White arrows - plasma membrane. Yellow arrows - electron-lucent layer in between the fibrillar primary cell wall and the PM. Corresponding angle from the tilt-series is shown on images. Scale bar = 200nm. Zoom-in field of view of 250nm.

### Differentiated Casparian strips

To demonstrate the power of our cryo-CLEM workflow, we decided to attempt the generation of cryo-lamellae by fluorescence marker-guided cryo-lift-out to obtain tomograms of Casparian strip (CS). CS begin to appear at about 800 µm from the root tip in a 5-day-old seedling with a root of 2-3 mm in length. Moreover, within the 80-120 µm thick root cylinder, CS are exclusively formed in a single root cell layer, the endodermis, and only occupy 2-3 µm thick region along the Z axis. Reliably positioning and identifying this region is only possible due to the CASP1-GFP marker, based on a strongly endodermis-expressed protein that strictly co-localizes with CS from their moment of inception^14^. Using our cryo-CLEM workflow, we always obtained cryo-lamellae containing the CS. Despite ice crystals formation in the vacuole, the CS and their associated PM appeared well-preserved (Fig. 3A, upper panel, movie 4). Again, cryo-ET delivered significantly greater details than resin embedded samples (Figure 3A, bottom panel). In TEM images of resin embedded samples, the CS appeared as a homogenous amorphous structure. Additionally, in these conditions, the CS can be identified as the region where plasmolysis-induced PM detachment is absent, due to the strong CASP-mediated wall adhesion of the CS membrane domain^15^. The PM at the CS moreover appeared to be more electron-dense than the rest of the PM (Figure 3A, bottom panel). By contrast, cryo-lamellae revealed a PM with repetitive structural elements, possibly representing the polymeric CASP scaffold and their associated proteins^14,16,17^ (Figure 3A, upper panel, movie 4). In addition, a membrane-proximal, electron-dense region, approximately four times the thickness of the PM itself was observed that might be due to the high density of cell wall enzymes associated with lignification during CS formation, such as peroxidases, dirigent proteins (DIRs) and others^17–19^. Finally, we were surprised to see that, despite a clearly increased contrast of the CS, certainly due to dense lignification of their cell wall (Fig. 3A, space between asterisks), the regularly spaced fibrillar arrangement of unmodified walls were still observable. This fits with similar observations in xylem vessels (see below). Cell wall areas with CS had to be imaged at lower beam intensity, because they were more sensitive to beam damage (“burned”) than other areas of the lamella. Since the same applies to xylem vessels, we assume that this enhanced sensitivity to the beam is due to the presence of lignin.

**Figure 3.**
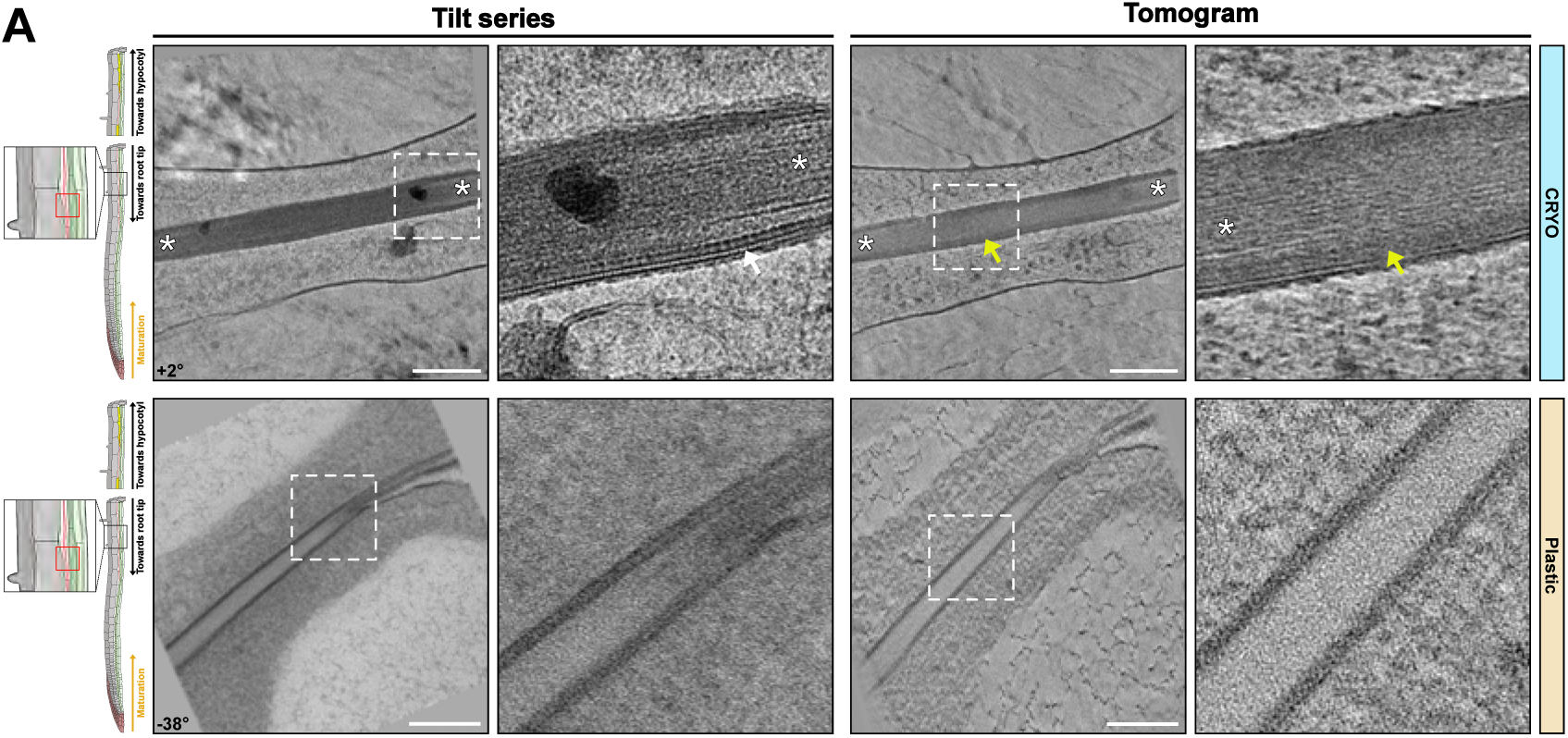
Tomograms of Casparian strip (CS). Example of cryo-fixed (upper panels, Movie 4) and chemically fixed, resin embedded conditions (bottom panels) of the established CS. White arrows – repetitive structural elements observed at the CS. Yellow arrows – membrane-proximal electron-dense region. White asterisk – unmodified cell wall. Corresponding angle from the tilt-series is shown on images. Scale bar = 200nm. Zoom-in field of view of 250nm.

### Nascent Casparian strips

Even more challenging than identifying CS for cryo-lift-outs is the generation of cryo-lamellae of the so-called string-of-pearls stage of initial Casparian strip formation^14^. The string-of-pearls stage occurs in a precise developmental window of about 1-2 cells among the more than hundred endodermal cells present in a 5-day-old seedling and again requires precise Z axis positioning within the root cylinder. Previous TEM analysis of this stage had revealed the presence of large numbers of extracellular vesiculo-tubular membranes^11^. The morphology of these extracellular membranes was extremely heterogenous probably due to the strong sensitivity of these structures to protoplast shrinkage during fixation (Fig. 4A-B, bottom panel). This contrasts with a more coherent shape and density of the extracellular membrane observed in cryo-conditions, with fields of vesiculo-tubular structures always residing in a lens shaped space between plasma membrane and the very thin, fibrous cell wall separating two endodermal cells. Tomograms of the string-of-pearls stage expectedly give divergent pictures, with either fully formed CS with little or no extracellular vesicles (Fig. 4C, upper panel, movie 7), an unformed CS with many vesicles (Fig. 4A upper panel, movie 5), or an intermediary stage with partially formed CS and fewer extracellular vesicles (Fig. 4B, upper panel, movie 6). This is consistent with current models according to which the string-of-pearls stage represents partially formed CS domains, with intercalated areas of strong secretion where the CS has not yet formed^20^. The ontogenesis of the extracellular vesicles during CS formation is unclear, but one straightforward explanation could be that they are the result of an excessive membrane surface that is resolved by extrusion of PM into an extracellular space between PM and primary cell wall. Indeed, vesiculo-tubular membrane structures tethered to the PM on its cytosolic face can easily be observed in this region (Fig. 4B, zoom-in). An intriguing question is the eventual fate of these extracellular membranes, since they are not present anymore upon CS completion (Fig. 4C, Fig. 3, movie 7, movie 4). Interestingly, we could observe extracellular membranes that appeared to become appressed, flattened against the primary cell wall, suggestive of turgor pressure-driven compression and eventual resorption of these structures back into the cell (Fig. 4A, upper panel, zoom-in). Very young CS regions reveal an even clearer nanostructure of the CS and its membrane domain (Fig. 4C, upper panel, movie 7). As described above, the PM often displays very strictly ordered repeats within the membrane (Fig. 4C, upper panel, zoom-in), associated with a more amorphous layer immediately outside of the PM, followed by a fibrous region of the wall, similar to unmodified primary wall, except for an increased width. Remarkably, no distinct middle lamella region can be observed within the primary cell wall. Despite resembling unmodified primary walls, the CS appeared always as slightly more electron-dense.

**Figure 4.**
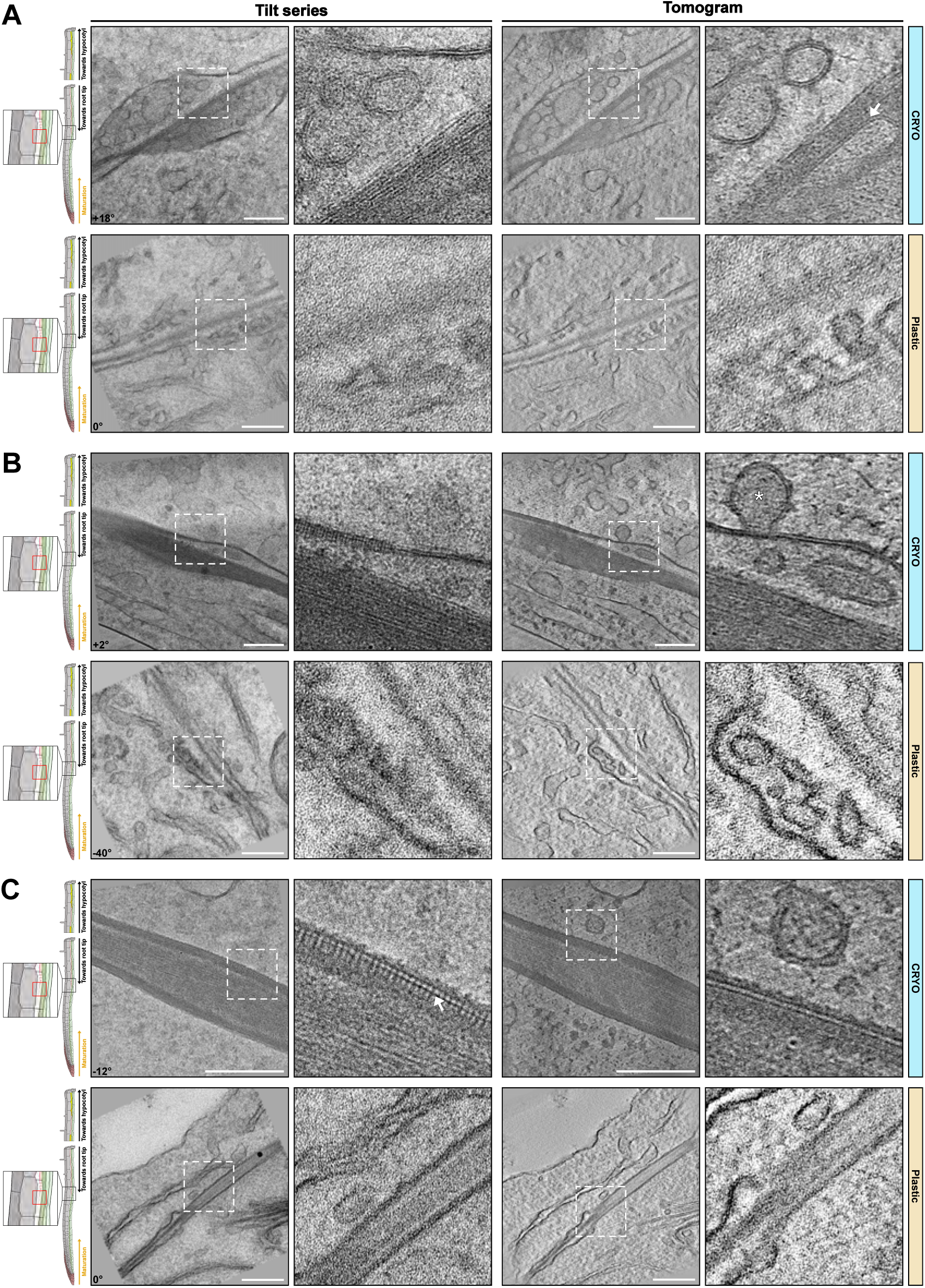
Tomograms of Nascent Casparian strip. Example of cryo-fixed (upper panels) and chemically fixed/resin embedded conditions (bottom panels) of nascent CS. **A**) Vesiculo-tubular structures in a lens shaped space between plasma membrane and the very thin, fibrous wall section between two endodermal cells (upper panels cryo). Similar structure in resin sections (bottom panels) showing tubular structures close to membranes but difficult to distinguish from artefactual membrane invaginations due to plasmolysis. White arrow - flattened vesicles against the primary cell wall. Movie 5. **B**) Intermediary stage with partially formed CS and fewer extracellular vesicles. White asterisk - vesiculo-tubular membrane structures tethered to the PM on its cytosolic side. Movie 6. **C**) Fully formed CS without extracellular vesicles (Movie 7). White arrow – nanostructures repeated within the membrane. Corresponding angle from the tilt-series is shown on images. Scale bar = 200nm. Zoom-in field of view of 250nm.

### Suberin lamellae formation

While also occurring exclusively in the endodermis within a primary root, suberin lamellae (SL) become deposited around the entire cell, in contrast to the very restricted formation of a CS band in the median position of endodermal cells. Suberin is chemically distinct from the lignin present in CS and occurs at a later stage of its development. It consists of hydroxylated fatty acids, which can become esterified to each other, but also to phenolics, such as ferulic acid, as well as glycerol. Despite a good knowledge of its monomeric composition, much remains unknown concerning its polymeric connections and the resulting spatial arrangements within the wall and how it is connected to unmodified or lignified primary cell wall^21^. Especially, there is still no widely accepted model that can explain the intriguing, lamellated appearance of suberin, as well as its positioning and integration between the cellulosic/lignified cell wall and the plasma membrane. We identified areas for cryo-lift-outs and lamellae preparation using a fluorescent reporter for the transcriptional activity of GLYCEROL-3-PHOSPHATE sn-2-ACYLTRANSFERASE 5 (GPAT5), a central and suberin-specific enzyme in the suberin biosynthetic pathway^22^. Suberin lamellae (SL) can be identified on tomograms of cell walls between an endodermal and its neighboring cortical cell, since only the cell wall on the endodermal side will display suberin formation. As described above, the unmodified cell walls showed a high degree of order with fibrous elements in parallel and equally spaced arrangements (Fig. 5A-D, upper panel, movie 8-11). We were at first confused to repeatedly observe the previously described precipitates, nicely aligned in the cell wall space between cortex and endodermis, but always “off-center”, i.e. closer to the endodermal side (Fig. 5A-D, movie 8-11). We now interpret this as the result of increased wall synthesis of the cortical cell as compared to the endodermal cell. This would place the middle lamella, the pectin and calcium-rich layer of attachment between plant cells, closer to the endodermal side. In the space between primary endodermal walls and plasma membrane, we could observe a thin layer of material, not present on the cortical side. Towards the PM, the suberin layer becomes wavy, or jagged, but remains separated from the plasma membrane by an additional, membrane-proximal layer of different density. We interpret the wavy/jagged aspect of the membrane-facing side of the suberin layer as partially-formed lamellae (Fig. 5A-D, yellow arrows). Indeed, close inspection of the suberin layer allows to identify two or three lamellae constituting the suberin layer at this stage. It is important to point out that this is the first time that the lamellated structure of suberin is visualized in the absence of any heavy metals as contrasting agents, demonstrating that there are indeed inherent structural differences detectable not only by heavy metal within the suberin layer but also by phase contrast used in cryo-EM. Moreover, the absence of fixation and dehydration/embedding procedures allows more reliable estimations of the thickness of the lamellae in our samples (5-7 nm, Fig 5B). Interestingly, when observed at the right tilt angle, suberin lamellae appeared of a contrast and thickness very close to that of the PM. This would fit with a model of a very regular polymeric arrangement in which one lamella could be defined by two units of hydroxy-fatty acids, parallelly aligned to each other, similar to fatty acids in the lipid bilayer of a membrane.

**Figure 5.**
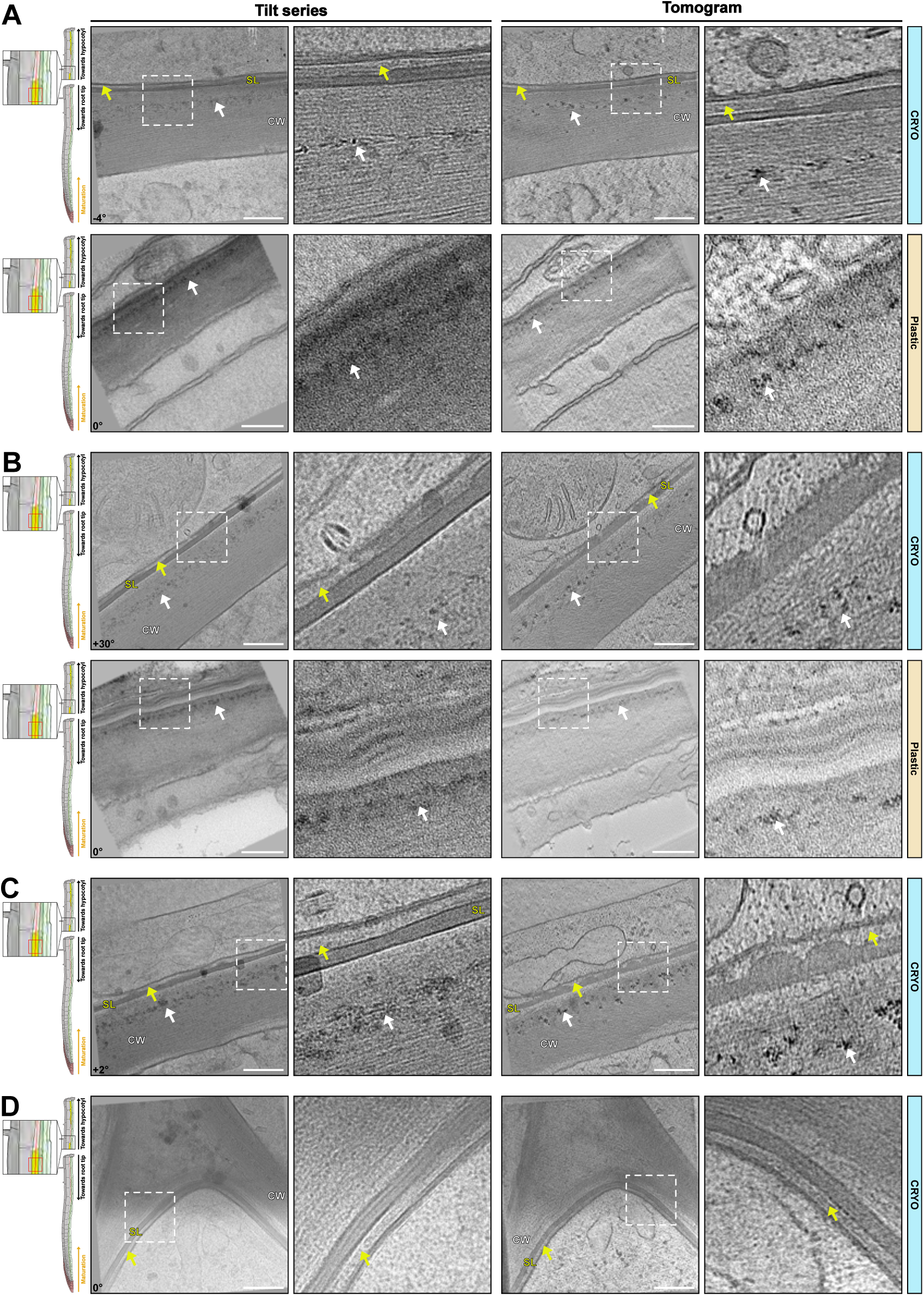
Tomograms of mature cell wall and early suberizing cells. **A-D**) Example of cryo-fixed (upper panels, Movie 8-11) and chemically fixed/resin embedded conditions (bottom panels) of the early suberizing cells. **A-C**) Tomograms of cell wall and early suberin deposition at the interface between endodermal and cortical cells. **D**) Tomograms of cell wall and early suberin deposition at endodermal cell corner. SL – suberin lamella. CW – cell wall. White arrows – electron-dense precipitates. Yellow arrows – electro-lucent layer in between PM and suberin lamellae. Corresponding angle from the tilt-series is shown on images. Scale bar = 200nm. Zoom-in field of view of 250nm.

### Xylem vessel cell walls

Areas containing the thick cell walls of xylem vessels can be identified by their autofluorescence for cryo-lift-outs. Xylem vessels are the cellular units crucial for water transporting vascular networks in plants and represent a type of heavily lignified cell that is very different from the endodermis. In contrast to the endodermis, lignification in xylem vessels is preceded by the formation of thick, secondary cell wall structures that are initially unmodified and rich in cellulose. This is then followed by strong lignification of the secondary wall thickening, in concert with the xylem cell undergoing programmed cell death, often leading to a boost in post-mortem lignification^23,24^. This is different than the endodermis, which stays alive during both lignification and suberization of its walls. Xylem vessels are devoid of cytoplasm and filled with xylem sap, containing dissolved minerals and comparatively little organic substances. We therefore expected heavy ice crystals formation but were surprised to see a very good degree of vitrification, both in the wall and in the vessel lumen. Again, highly organized fibrillar structures were observed in the xylem secondary walls, despite the presence of lignin, indicated by the higher electron-density of its cell wall compared to those of the neighboring cell (Fig. 6A-B, L indicates denser, lignified wall region). The neighboring cell, whose intact cytoplasm would indicate it to be a xylem parenchyma cell (Fig. 6A-B, movie 12-13), displayed a much more disordered arrangement of microfibrils.

**Figure 6.**
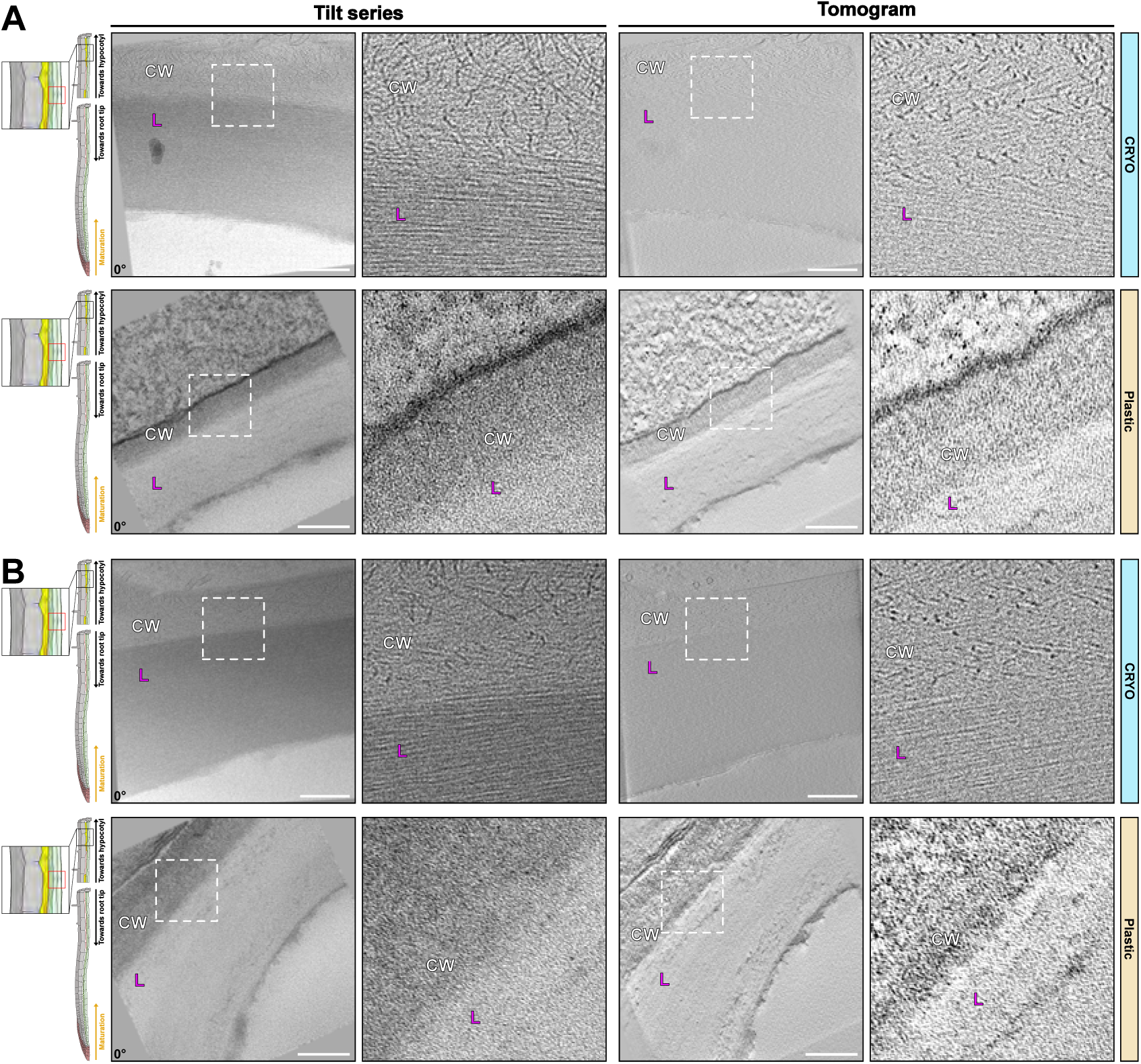
Tomograms of lignin-impregnated cell wall of xylem vessels. **A,B**) Example of Cryo fixed (upper panels, Movie 12-13) and chemically fixed/resin embedded conditions (bottom panels) of Xylem vessels. CW – Cell wall of non-lignified cells. Magenta L – lignin impregnated cell wall from xylem cells. Corresponding angle from the tilt-series is shown on images. Scale bar = 200nm. Zoom-in field of view of 250nm.

Nothing resembling these fibrillar structures could be observed in comparable images of resin embedded samples. Resin sections revealed the difference between xylem vessel and parenchyma walls through a higher degree of staining of the parenchyma cells, which consequently appeared more electron-dense than the lignified xylem vessel walls. In the cryo-lamellae, the differences in wall structure were already apparent from the very different orientation and degree of order of the fibrillar wall component (6A and B, zoom-in).

### The Cryo-Lift-Out Process

The cryo-lift-out process typically involves several key steps. Here, we will focus on the main challenges, and solutions we developed in our attempts to initially target the Casparian strips (CS). The complete workflow used to reach proper targeting will be presented. CS were visualized using the CASP1-GFP marker under its endogenous promoter^25^. Roots of the transgenic lines were rapidly frozen using HPF to preserve its native structure. After having tried different protocols, the best freezing results, minimizing ice crystals formation and cell osmotic perturbation, were obtained by cutting the tip of *A. thaliana* root, transfer it rapidly to 20% dextran inside a 100 μm deep carrier of 3mm diameter and immediatly high-pressure freeze it (Fig. 7A).

**Figure 7.**
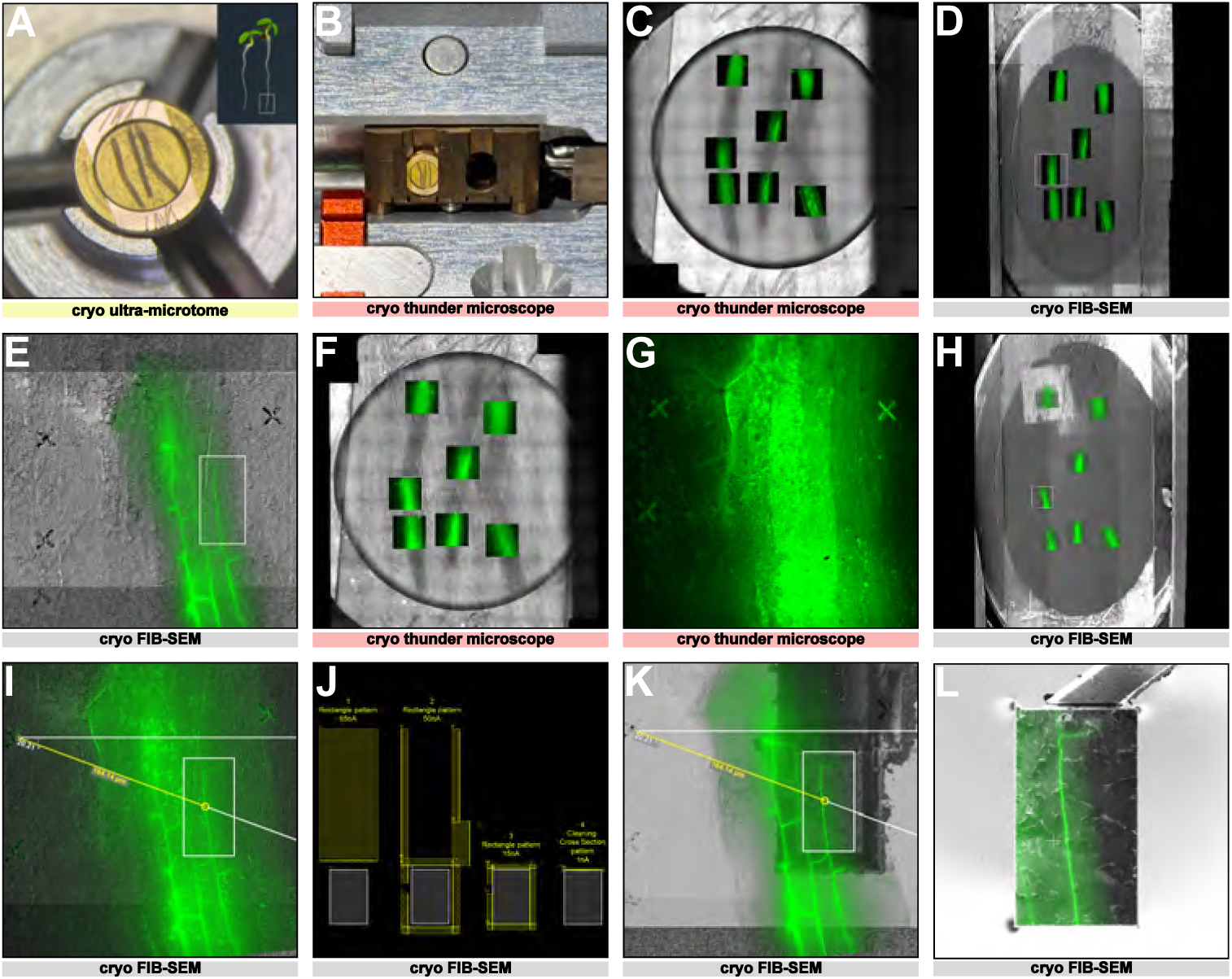
Coarse and fine targeting using fluorescence Z-stacks and fiducials marks. **A)** Carrier containing three root segments in the cryo-ultramicrotome chamber. The carrier surface is polished with a diamond knife and fiducial marks (scratches) are engraved on the carrier rim using a diamond tip. **B)** Carrier mounted in a modified cryo-cassette for imaging in the Leica Thunder cryo-LM. **C)** Overview map of the carrier in reflective light showing the localization of the fluorescent Z-stacks of the CASP-GFP signal acquired in the cryo-LM. **D)** Full map of the carrier acquired with the ion beam in the cryo-FIBSEM using MAPS software (TFS) overlaid with the fluorescent Z-stacks. **E)** Enlarged view of the white square in D showing fiducials crosses milled at 100 µm of distance around the ROI. **F)** Reflected light overview map of the carrier showing the fluorescent Z-stacks from the second round of cryo-LM with milled fiducials crosses. **G)** Enlarged view of the white square in F with the crosses at the surface of the carrier. **H)** Full map of the carrier acquired with the ion beam in the cryo-FIBSEM overlaid and accurately aligned using the fiducials crosses of the fluorescent Z-stacks. **I)** Enlarged view of the white square seen in H showing a maximum intensity Z projection containing the crosses and the target aligned with the ion beam snapshot. Polar coordinates measure the angle and distance of the target from one cross. **J)** Four sequential milling patterns (yellow) are used to prepare the block (55 x 80 µm white rectangle). **K)** Block prepared with the ion beam, overlaid with the fluorescent target. The block preparation includes a long opening trench at 65 nA, trenches around the block at 50 nA and then at 15 nA, undercut at 1 nA with 10° milling angle and needle attachment face polishing at 1 nA. **L)** Block attached to the silver needle through silver redeposition overlaid with the fluorescent target. The block width is reduced to 40 μm in width and the bottom is polished at 5 nA.

### Targeting the Casparian strip with a precision of approximately 1 µm in XY and Z by a 2-step targeting process

Once the root tip is vitrified in the carrier, the first step is to trim the surface by using a cryo-ultramicrotome equipped with a cryo-trim knife. This allows us to reach the appropriate distance in Z of the region of interest (ROI) as well as to smooth the vitrified sample surface for visualization of the fluorescent signal. The carrier is then transferred to a cryo-Light Microscopy (cryo-LM) equipped with a cryo-stage. To insert our sample into the cryo-LM, the holder has been modified to accommodate the carrier (Fig. 7B). The entire surface of the carrier is imaged in the fluorescence channels, as well as reflected light, creating a map of the whole surface (Fig. 7C). Z stacks of the different ROI are also obtained at a resolution of 1µm Z steps to determine where the cryo-lift-out block will be carved.

The vitrified sample is then transferred to the cryo-FIBSEM system. The FIB can be precisely controlled to mill specific ROI. In the case of the root tip, images of the entire surface obtained by Cryo-LM are transferred in the software of the cryo-FIBSEM and used as a map to identify the different ROI. Since the cryo-FIBSEM used in these experiments is not equipped with an in-chamber light microscope, the targeting relies only on the perfect overlay of fluorescent maps on the cryo-FIBSEM images and precise measurements.

To improve the CS targeting, we developed a routine in two steps. First, the cryo-LM map of the entire surface of the carrier is overlaid with a map of the entire surface of the carrier obtained by scanning with the ion beam. Then, the two maps are overlaid and the root tips identified (Fig. 7D). Three small crosses acting as fiducial marks are carved with the FIB at the surface of the sample surrounding ROIs (Fig. 7E). Once done, the carrier is brought back to the cryo-LM and precise Z stacks containing fiducial marks are acquired for each ROIs (Fig. 7F-G, Movie 14). Then, the carrier is reloaded into the cryo-FIBSEM. The precise distances in X, Y and Z between the ROI and each of the fiducial marks are measured on the fluorescent images (Fig. 7H-I) and reported on the cryo-FIBSEM images (Fig. 7K). The fiducial marks are used to determine the size and depth of the block that will be prepared for cryo-lift-out (Fig. 7J). The typical size of a block is 50 x 80 µm and 30 µm in depth (Fig. 7L).

### Extended Operation Through Enhanced Cooling System Design

To enable extended cryo-FIBSEM operations, we developed a copper spiral cold trap that integrates into the existing microscope cooling system (Fig. 8). The trap consists of a manually coiled copper tube, providing a larger inner diameter (4 mm) and an additional cooling line length (70 cm) (Fig. 8A). The larger diameter of the copper tube prevents clogging while maintaining efficient cooling. The trap is connected to the standard cooling lines (Fig. 8B). The design enables compact integration within the microscope’s heat exchanger housing without compromising accessibility for refilling the cooling dewar (Fig. 8C). Performance tests demonstrated that the spiral cold trap extends continuous operation from the previous 12–24 hours limit to over 5 days without system clogging. This enhancement enables completion of entire sessions without interruption. The system maintains stable cryo-conditions eliminating the risk of ice contamination of the sample during transfer and thermal cycling of the microscope during multi-day sessions. The cooling dewar is refilled twice per day.

**Figure 8.**
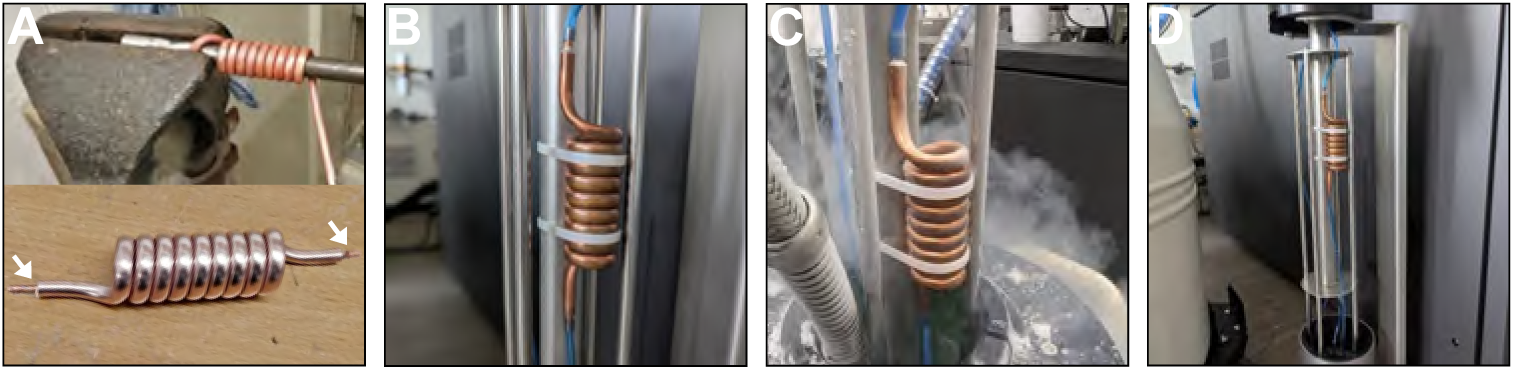
Design and implementation of a copper spiral cold trap. **A) Top,** Fabrication of the cold trap showing manual coiling of a 70 cm long copper tube (6 mm outer diameter, 4 mm inner diameter) around a 16 mm steel bar to create the spiral geometry. **Bottom,** Two tubes (arrow), 1 cm long and 4 mm outer diameter are soldered at both ends for connection into the cooling system (arrow). **B)** Installation of the cold trap in the microscope’s heat exchanger, showing connection with the existing blue cooling lines. **C)** The heat exchanger lifted for LN_2_ refilling. **D)** Wide view of the complete cold trap assembly mounted in the heat exchanger housing.

### Silver-Plated Needles for Direct Sample Manipulation

To address the time-consuming nature of adaptor chunk preparation^8,9^, we developed a silver-plated needle system enabling direct sample attachment. The needles are prepared through electroplating of standard tungsten EasyLift^TM^ needles in silver cyanide bath (Fig. 9A). Two fine-silver anodes and precise control of voltage and current ensure homogeneous plating. The plated needles undergo a systematic preparation protocol utilizing the EasyLift^TM^ rod’s rotational capability (Fig. 9B). We verified that 90° clockwise and counterclockwise rotations could be performed under both, ambient and cryo-conditions without compromising vacuum and needle temperature integrity. This rotation enables precise shaping of the needle through a series of milling steps (Fig. 9C). The final needle measure 250 µm in length and thinned to a final thickness of 17 μm in both dimensions, giving a 25 μm horizontal attachment surface. While initial preparation requires approximately 4 hours, the resulting needle can be used for months of operation.

**Figure 9.**
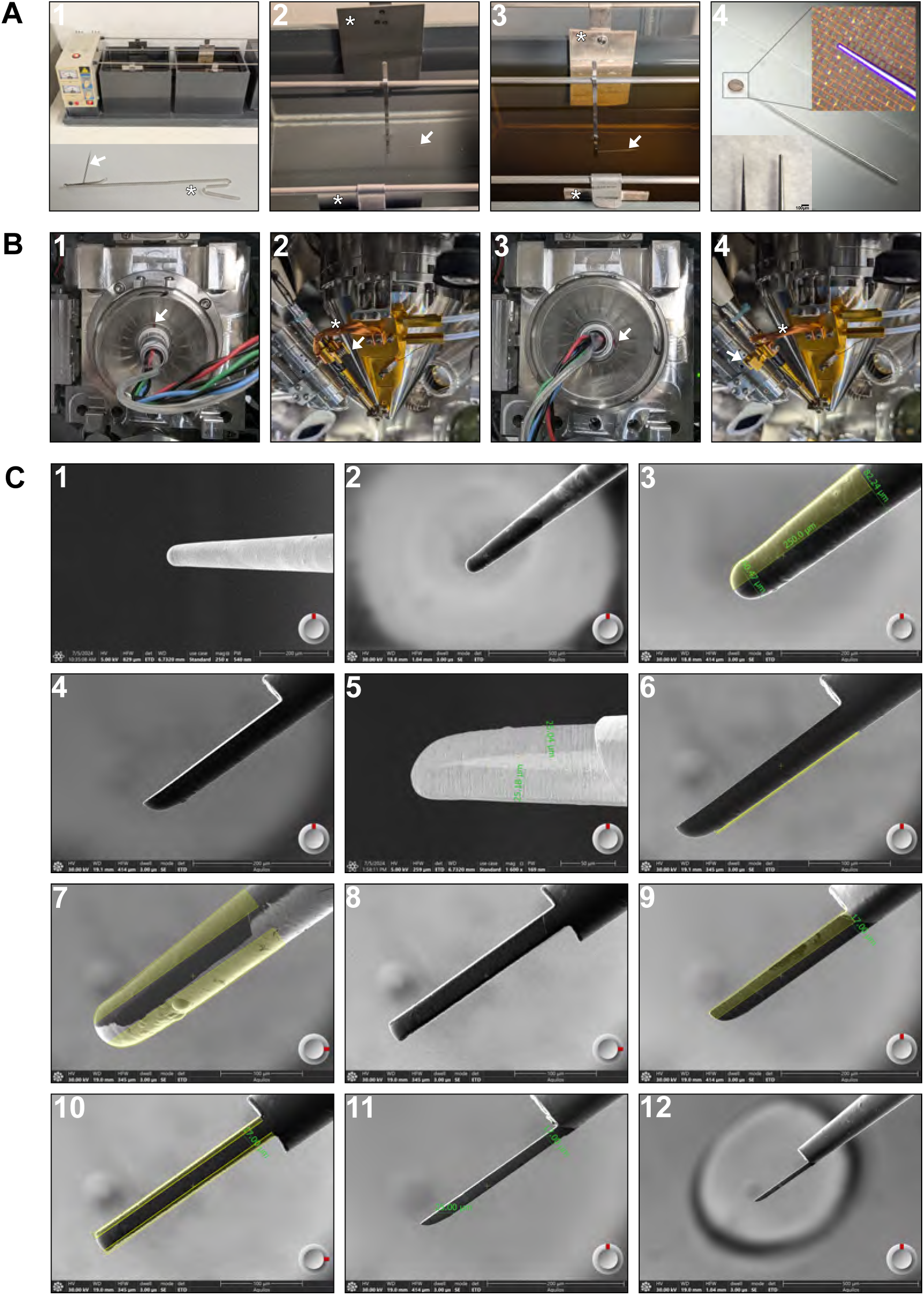
Preparation of the silver coated EasyLift^TM^ needle. **A)** Electroplating setup for silver coating of tungsten EasyLiftTM needles**. 1**) Complete electroplating apparatus showing the rectifier on the left, an electrolytic degreasing bath and an electrolytic silver bath with two fine silver anodes (upper panel). Needle holder (asterisk) made with a silver-plated copper strip (3mm width, 0.5mm thickness) maintaining the tungsten needle (arrow). **2**) Close-up view of the degreasing bath with the two stainless steel anodes (asterisks) and the needle attached to the needle holder (arrow). **3**) Close-up view of the silver bath with the two silver anodes (asterisks) and the needle (arrow). **4**) Silver needle tip thickness estimation under a binocular using a 300 square mesh copper grid attached with a transparent tape to a glass slide. Comparison of the original tungsten needle (bottom left) and the silver plated needle (bottom right) (scalebar 100µm). **B)** Rotation of the EasyLift^TM^ rod. **1**) The EasyLift^TM^ rod in the standard position, seen from outside the Aquilos^TM^ 2 chamber. The red mark of the connector position is upward (white arrowhead). **2**) The EasyLift^TM^ seen from inside the microscope, showing the clamp (white arrowhead) of the cooling braid (asterisk) placed to the right of the EasyLift^TM^ end. **3**) The same view as (1), but rotated 90° CW. The red mark is now on the right, at 90° compared to the previous position. **4**) The view from inside the microscope with the 90° CW rotation, showing the position of the cooling braid clamp is now downward. **C**) Milling steps for preparing the silver needle**. 1-2**) Initial views of the silver-plated needle in electron beam (EB) and ion beam (IB) views. The bottom right circle with the red mark indicates the EasyLift^TM^ rotation. **3-4**) A polygonal pattern is drawn at the needle tip to remove the top half of the needle material. **5**) EB view reveals the silver layer thickness around the tungsten core. **6**) A thin milling pattern is drawn to create a flat bottom face. **7-8**) EasyLift^TM^ rod rotation of 90° CW (red mark to right) exposes the bottom face. Two patterns are drawn to thin down the needle to 25μm. **9**) EasyLift^TM^ rod rotation 90° CCW to original position (red mark up). A milling pattern is drawn to remove remaining tungsten and achieve 17 μm thickness. **10**) EasyLift^TM^ rod rotation 90° CW enables thinning to 17 μm in the second dimension. **11-12**) Final 90° CCW rotation returns needle to the original position. The 17 μm thickness gives a needle width of 25 μm in the horizontal dimension, providing sufficient surface area for robust sample attachment.

### Enhanced Sample Attachment Performance

The silver needle enables direct attachment to the sample blocks without intermediate adaptor chunks (Fig. 10). Silver’s higher sputter rate compared to copper facilitates robust attachment through single-pass regular cross-section milling. We demonstrated successful cryo-lift-out and attachment of sample lamellae between grid bars (Serial Lift-Out, Fig. 10A) or onto silver film supports (SOLIST, Fig. 10B). This approach eliminates the 23 minutes chunk preparation previously required and reduces total preparation time per sample block. The simplified workflow involves a single welding step rather than the traditional dual welding process, decreasing the risk of sample loss during cryo-lift-out and serial sectioning (Movie 15-16).

**Figure 10.**
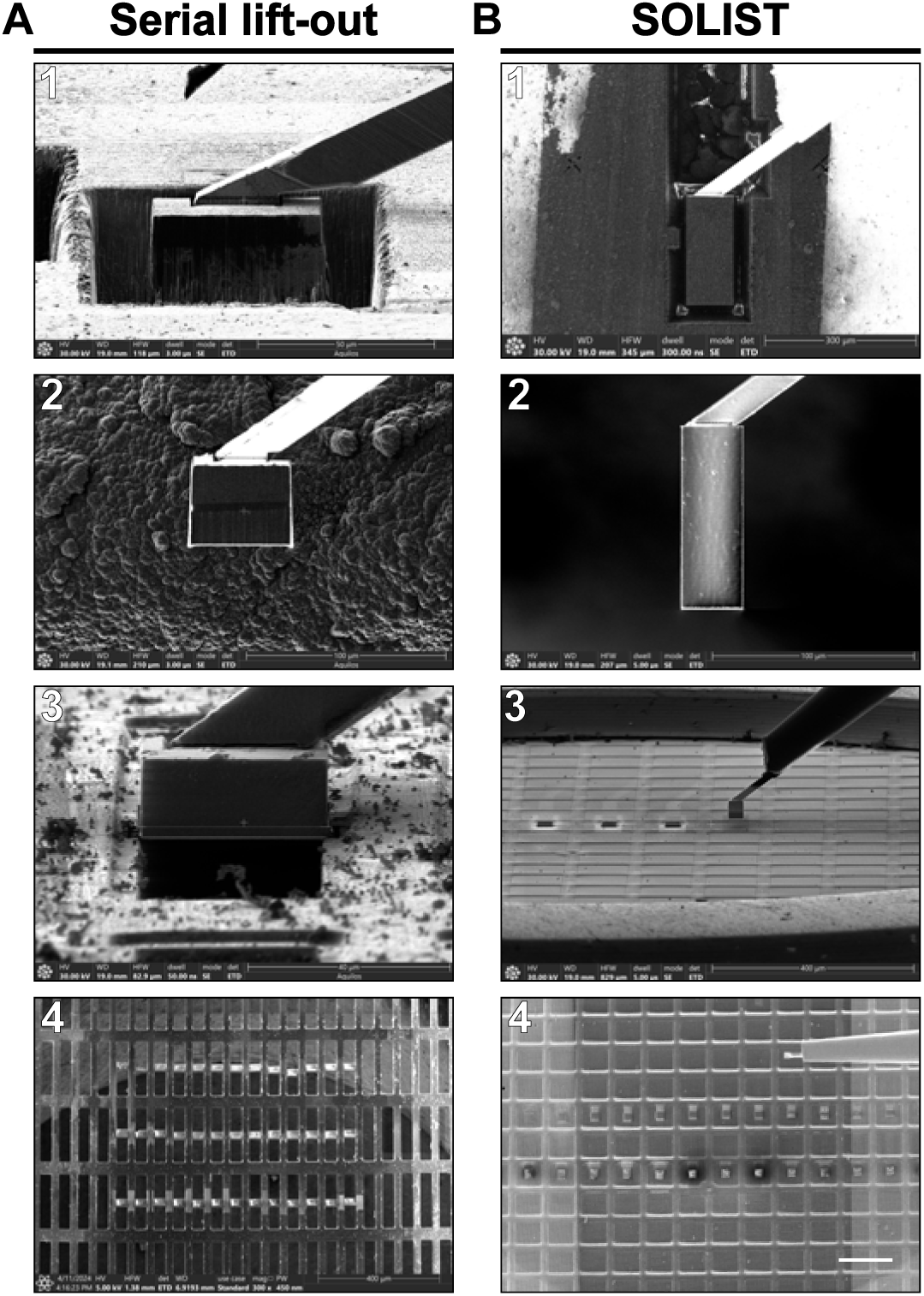
Direct sample attachment using silver needle. **A**) Serial Lift-Out method. **1**) Direct attachment of silver needle to the sample block from a high-pressure carrier using single-pass regular cross-section milling. **2**) Lifted-out sample block. **3**) Double-sided attachment between grid bar of a 100-400 rectangular mesh copper grid. **4**) Full grid view showing lamellae before final polishing. **B)** SOLIST method. **1**) Direct attachment of silver needle to the sample block from a HPF carrier using single-pass regular cross-section milling. **2**) Lifted-out sample block. **3**) Sample block lift-out after successful lamella deposition on silver support film. **4**) Image of receiver grid showing lamellae before final polishing (scale bar 200 µm).

### Enhancement of lamella stability through attachment to silver-coated support films

As a receiver grid for SOLIST lamellae we used standard 200 mesh copper R2/2 Quantifoil grid sputter coated with silver to obtain a film thickness of 500 nm. The thick metal layer enabled the attachment of the lamellae using single pass regular cross section pattern by redeposition of silver on the two sides of the lamella (Fig. 11). The attachment strength was tested by pushing the lamella using the silver needle with 10 steps of 200 nm (Movie 17). This strong attachment and the film thickness make a stable system which enables very stable polishing steps. We can typically polish 20 µm wide lamellae without lamellae bending, up to a thickness of 150-200 nm

**Figure 11.**
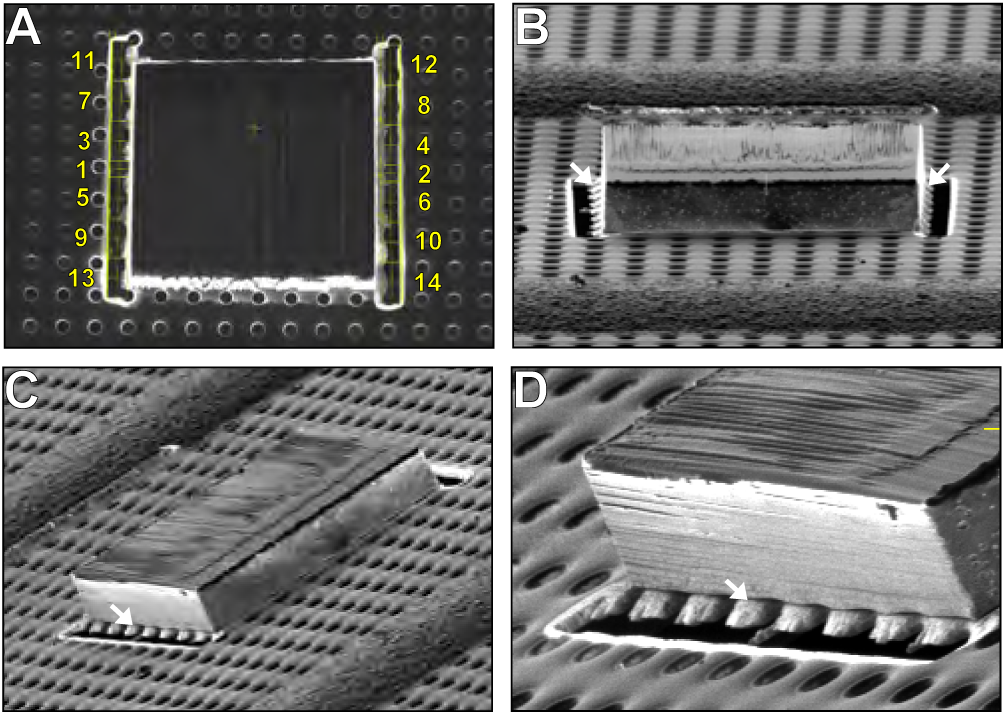
Lamella attachment on silver film. **A**) Lamella (width 30 µm, height 27 µm, thickness 5 µm) perpendicular to the ion beam showing single-pass regular cross-section milling patterns (yellow rectangles) for attachment on both sides of the lamella. The numbers from 1 to 14 indicate the milling sequence. **B)** Lamella at 10 degrees to the ion beam showing the silver redeposition on both sides of the lamella (white arrows) making a strong attachment to the silver film. **C)** The same lamella rotated (R -110°) showing the attachment on the side of the lamella (white arrow). **D**) Enlarged view of the silver redeposition (white arrow) between the silver film and the lamella.

### Lamella preparation for curtaining reduction

The curtaining propensity during focused ion beam milling is predominantly influenced by the surface topography of the milling front. While chemical vapor deposition of organometallic platinum via gas injection system (GIS) can help planarize the surface prior to milling, its effectiveness is contingent upon the initial smoothness of the substrate. As demonstrated in Fig. 12B, organometallic platinum deposition on a roughly milled surface results in inhomogeneous protective layers with localized discontinuities that are readily apparent in cryo-TEM imaging (Fig. 12B 2). These heterogeneities in the protective layer contribute significantly to differential milling and subsequent curtaining artifacts in the final lamella. Although the conventional GIS angle of incidence (30°) may contribute to incomplete filling of surface features, even perpendicular GIS deposition does not fully eliminate these discontinuities (Fig. 12B). Through systematic optimization, we established a protocol that substantially reduces curtaining. Notably, low-current polishing of the milling front without initial thin organometallic platinum deposition on the top of the lamella proves insufficient for curtaining reduction. The sequential application of a thin organometallic platinum layer on the top of the lamella, followed by a front surface polishing, and a second organometallic platinum deposition on the front of the polished lamella minimizes curtaining artifacts (Fig. 12A 1-3). The enhanced control over lamella geometry afforded by the serial lift-out method enables precise manipulation of the milling front topography, thereby optimizing protective layer uniformity and ultimately reducing curtaining in the final lamella (Fig.12A 4). This approach represents a significant improvement providing a reproducible method for high-quality lamella preparation with minimal curtaining artifacts (Fig. 13).

**Figure 12.**
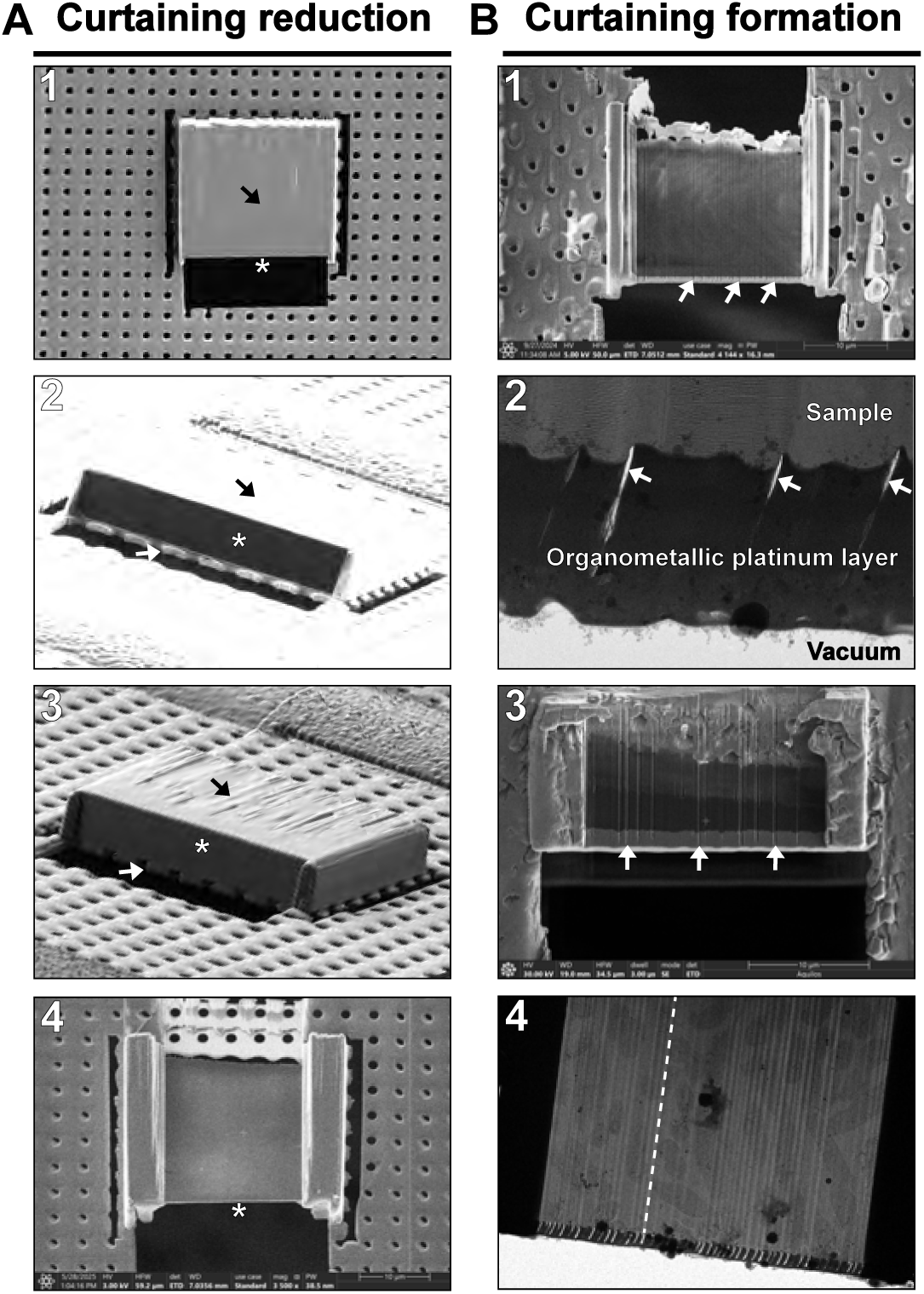
Curtaining reduction strategy. **A)** Optimized lamella preparation. **1**) Same lamella seen in Figure 11, perpendicular to the ion beam. Front surface polishing (asterisk) at high current (1 nA) after initial organometallic platinum deposition (20 seconds) by gas injection system (GIS) perpendicular to the top surface of the lamella (black arrow). **2**) Lamella at 10 degrees to the ion beam and rotated (R -30°) showing the lamella front surface (asterisk) after polishing at 1 nA (the high contrast allows the visualization). The black arrow indicates the top surface of the lamella with the organometallic platinum layer, and the white arrow indicates the thick silver film. **3**) Lamella seen after organometallic platinum deposition on the polished front surface (asterisk). **4**) Optimized prepared lamella exhibiting a continuous, void-free organometallic platinum protective layer and exceptionally smooth lamella surface without curtaining artifacts after polishing steps. Imaged with the electron beam at 3 kV showing very low charging effect on the lamella. A very thin organometallic platinum layer remains at the front of the lamella (asterisk). **B**) Curtaining formation during lamella preparation. **1**) Lamella showing a very high curtaining effect after low-current (50 pA) polishing of the milling front without initial thin organometallic platinum deposition on top of the lamella. The organometallic platinum layer shows a high number of cavities (white arrows). **2**) TEM view of the same lamella revealing void formation within the organometallic platinum layer (white arrows). These elongated cavities are consistently inclined at approximately 30° to the lamella surface, directly corresponding to the incident angle of the GIS precursor flux. This void distribution suggests shadowing effects during the deposition process. **3**) Organometallic platinum deposition performed perpendicular to the lamella front (achieved by rotating the stage to 30°, aligned to the GIS column) after low-current polishing. Despite the modified deposition geometry, similar voids persist in the protective layer (white arrows), indicating that deposition angle alone is insufficient to eliminate porosity in the organometallic platinum layer and its presence is not related to a shadowing effect. **4**) TEM lamella overview showing the very high curtaining effect (parallel to the white dash line) due to the cavities in the organometallic platinum layer.

**Figure 13.**
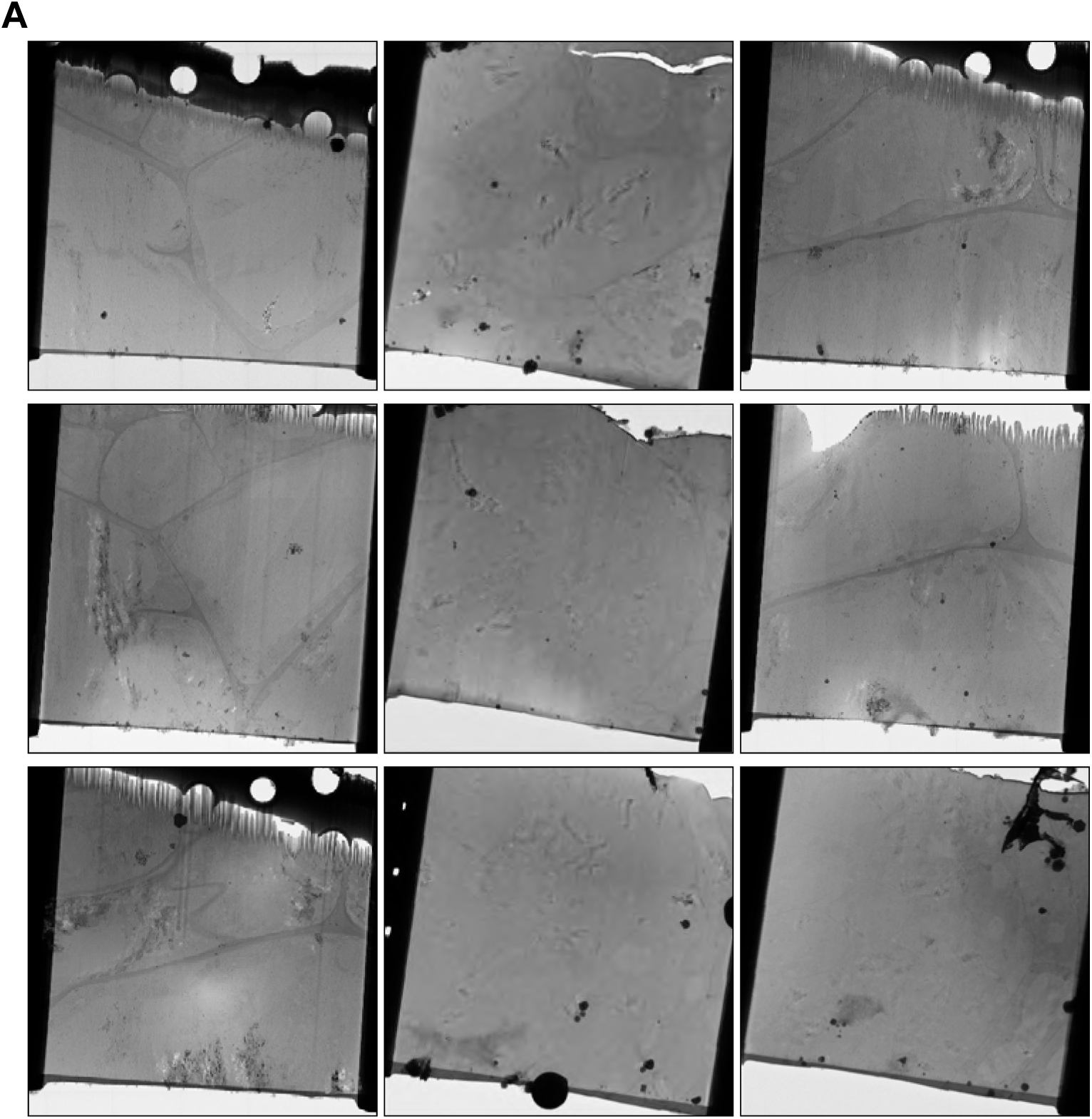
Examples of cryo-lamellae overviews in cryo-TEM. Lamellae followed the reproducible curtaining reduction strategy described above. Lamellae width is 20 µm, height can vary.

## Discussion

Unlike animals, plants have evolved complex multi-cellularity based on walled, pressurized cells, allowing plants to rapidly grow and maintain cellular structure by generating large internal vacuoles and high osmotic pressure. This internal pressure generates turgidity, because it is resisted by intricately structured cell walls. Cell wall nanostructure is dynamically modified by plant cells during growth, development and adaptation to environmental challenges. A plethora of cell-type-specific cell wall modifications are crucial for allowing many plant cells to carry out their specific function. Cell walls provide structure and coherence to plant tissues and have rendered them a favorite object of investigation since the beginning of microscopy^26^. However, these same features have also rendered plant tissues a challenge for cryo-ET, since large vacuoles, pressurized cells and rigid cell walls, impede efficient vitrification. Vacuoles increase the likelihood of ice crystals formation, which can severely compromise sample integrity and resolution. Furthermore, the high internal pressure and associated rigidity of the plant cell also limits the tolerance of the tissue to mechanical constraints, especially osmotic stress, which can rapidly induce severe artifacts, especially at the crucial plasma membrane-cell wall interface. Therefore, while it is commonly accepted to add cryoprotectant to improve vitrification of animal samples, we use cryoprotectant only to fill the carrier some seconds before freezing to avoid osmotic artefacts. Impermeability of cell walls can also reduce the even penetration of cryoprotectants, complicating efforts to achieve complete vitrification. While complete vitrification remains a technical hurdle that needs to be solved, our protocol demonstrates significant improvements in sample handling and structural preservation, making previously inaccessible plant tissues amenable to cryo-ET analysis. Despite these challenges, we successfully imaged Casparian strips, as well as suberized cell walls in cryo-lamellae, demonstrating the power of our enhanced cryo-CLEM workflow. Casparian strips are highly localized, ring-like impregnations of primary cell walls with hydrophobic lignin, generating a critical, extracellular diffusion barrier across the endodermis, allowing the root to regulate solute and water transport. Later, endodermal cells further suberize, forming so-called suberin lamellae between the plasma membrane and primary cell wall on all endodermal surfaces, fully sealing off the endodermal cell from its environment with this hydrophobic, cork-like secondary cell wall. It was indeed the observation of a field of highly suberized, dead cells that Robert Hooke observed in sections of cork tissue and which led to his coinage of the term “cells”^26^. Our work now visualizes endodermal suberin and lignin structures in near-native conditions and allow an unprecedented examination of their architecture and formation. Our data reveal an unexpectedly high order and spatial regulation of fibrils in the cell wall, with surprising degrees of parallel arrangements distal to the plasma membrane and more disorganized arrays closer to the PM. While the CS PM domain has previously been reported to be more electron dense and to display tight attachment to the plasma membrane, cryo-ET now reveals the CS-associated plasma membrane to form a much thicker, electron-dense layer than previously observed, possibly representing the dense accumulation of extracellular and transmembrane proteins known to localize to this domain. The unmodified primary cell wall of endodermal cells, consisting of cellulose microfibrils, hemicelluloses and pectins, also display a degree of parallel order that we do not observe in conventional TEM. In cell corners a much more disorganized arrangement of fibrils is observed. Furthermore, cryo-ET of suberin *in situ* provide the first visualization of lamellated suberin in the absence of any EM staining and embedding that could have altered their native structure. This demonstrates that suberin lamellae are indeed made of polymers of different electron densities and do not simply have differing affinities for EM stainings.

Our improvements in cryo-lift-out techniques—including the use of silver-plated EasyLift^TM^ needles and an extended cryo-FIBSEM operational workflow—have improved targeting, while insuring the generation of many lamellae of high quality from a region of interest. These advancements significantly enhance the reproducibility and feasibility of cryo-ET for many previously challenging or inaccessible tissue or organs.

While our method relies on automation software such as AutoTEM^TM^ Cryo for lamella preparation and polishing, a significant portion of the workflow remains manual. Critical steps including block preparation for lift-out, lamella attachment to grids, front-face polishing, and organometallic platinum deposition are currently not supported by automation software and therefore done manually. Development of softwares allowing automation of such tasks is of great importance to improve and facilitate cryo-FIBSEM lamellae generation.

In summary, while vitrification remains a persistent challenge, the findings presented here represent a significant advancement in cryo-electron microscopy. Further refinements of vitrification protocols and cryo-ET methodologies will continue to push the boundaries of structural plant biology, offering deeper insights into the fundamental processes governing root function and environmental adaptation.

## Methods

### Room temperature TEM analysis

Samples were prepared as described previously^11^. Plants were fixed in glutaraldehyde solution (EMS, Hatfield, PA) 2.5% in phosphate buffer (PB 0.1 M [pH 7.4]) for 1 h at RT and postfixed in a fresh mixture of osmium tetroxide 1% (EMS, Hatfield, PA) with 1.5% of potassium ferrocyanide (Sigma, St. Louis, MO) in PB buffer for 1h at RT. The samples were then washed twice in distilled water and dehydrated in ethanol solution (Sigma, St Louis, MO, US) at increased concentrations (30%–40 min; 50%–40 min; 70%–40 min; 100%–2x 1 h). This was followed by infiltration in Spurr resin (EMS, Hatfield, PA, US) at graded concentrations (Spurr 33% in ethanol - 4 h; Spurr 66% in ethanol - 4 h; Spurr 100%–2x 8 h) and finally polymerized for 48 h at 60 °C in an oven. For electron tomography, semi-thin sections of 250 nm thickness were cut. The area of interest was taken with a transmission electron microscope JEOL JEM-2100Plus (JEOL Ltd., Akishima, Tokyo, Japan) at an acceleration voltage of 200 kV with a TVIPS TemCamXF416 digital camera (TVIPS GmbH, Germany). Micrographs were taken as single tilt series over a range of −60° to +60° using SerialEM^27^ at tilt angle increment of 1°. Tomogram reconstruction was done with IMOD software^28^.

### High Pressure Freezing

Root segments (2 mm) from Arabidopsis thaliana were placed in 100 µm deep cavities of 3 mm gold-coated copper carriers (Type A, Wohlwend GmbH, Switzerland) filled with 20% (w/v) dextran (Sigma, St. Louis, MO) dissolved in MOPS buffer. The carrier was covered with a Type B carrier coated with hexadecene (Sigma, St. Louis, MO) and vitrified using a Leica EM ICE high-pressure freezer (Leica Microsystem, Vienna, Austria). After freezing, carriers were stored in cryobox in liquid nitrogen until further processing.

### Cryo-ultramicrotomy

The carrier was transferred to a Leica UC6 ultramicrotome (Leica Microsystem, Vienna, Austria) cooled to -170°C and the surface was trimmed using a Trim20 diamond knife (Diatome, Biel, Switzerland). After complete polishing of the surface, fiducial marks (scratches) were engraved on the carrier rim using a diamond tip (EMS, Hatfield, PA, US) to facilitate later alignment and correlation (Fig. 7A). The carrier was then stored in a cryobox in liquid nitrogen.

### Cryo-correlative light and electron microscopy workflow

For the first acquisition in the cryo-LM, the carrier was mounted in a modified cryo-cassette (Fig. 7B) for the Leica Thunder cryo-LM (Leica Microsystem, Vienna, Austria) and transferred to the cryostage for imaging. A complete overview map (11x12 tiles) of the carrier was acquired using reflected light with a 50x objective (NA 0.75, pixel size 260nm), using four focus points set to ensure sharp focus across the entire surface. Fluorescence z-stacks of the CASP1-GFP signal were collected using a GFP filter set (EX: 450-490, DC 495, EM: 500-550). The Z-stacks were initiated at the carrier surface and extended below the cells of interest with a step size of 1 μm (Fig. 7C). The carrier attached to the cryo-cassette was mounted on a 45° pretilt shuttle, gently cleaned with a fine brush if ice contamination was present and transferred to the Aquilos 2 cryo-FIBSEM (Thermo Fisher Scientific Inc., US). Remnant of ice contamination was removed quickly using a 5nA ion beam at fast scan (50ns, 768x512) at milling position. The carrier was then sputter coated with platinum for 15 seconds. The center of the carrier was set to eucentric height, rotated to 110° and tilted to 7°, perpendicular to the ion beam. A full carrier montage was acquired using the ion beam in MAPS software (Thermo Fisher Scientific Inc., US), with 6×11 tiles, overlap of 10% in X and 50% in Y, tile HFW of 950µm, 1536x1024, dwell of 1µs, 4 frames, and three focus points ensuring sharp focus of the carrier surface. The resulting ion beam map exhibited after stitching, an elongation along the vertical axis, a known limitation of MAPS software when using ion beam imaging. Fluorescence data was imported into MAPS and aligned to the ion beam image using three reference points (Fig. 7D). Despite the image distortion along the vertical axis, the overlay enabled direct stage navigation to regions of interest (ROIs). Because each montage has small local misalignments due to the stitching process, aligning the two montages together can result in a mislocalisation of the target up to 15-20 µm. To overcome this problem, three cross-shaped fiducial markers were milled at 100 μm of distance from each ROI using a 1nA ion beam current (crosses: 2 x rectangular pattern, X 20µm, Y 2µm, Z 0.2µm) (Fig. 7E). This distance should be sufficient to avoid damage to the sample while keeping the fiducials within the field of view of the cryo-LM. Once all the targets were surrounded by fiducial crosses, the carrier was then transferred back to the Leica Thunder cryo-LM. The carrier was imaged again as described above, but the acquired Z-stacks including now the milled fiducial crosses (Fig. 7F-G). These images would serve as the foundation for precise three-dimensional targeting of the regions of interest. Following imaging, the carrier was removed from the modified cassette and stored in a cryobox in liquid nitrogen.

### Silver support film for SOLIST lamella deposition

For SOLIST implementation, we used standard R2/2 Quantifoil carbon film on 200 mesh copper grids (Quantifoil Micro Tools GmbH, Germany) sputter-coated with 500 nm of silver using a Safematic CCU-010 sputter coater (Safematic GmbH, Switzerland) at an argon pressure of 5x10-2 mbar and a process current of 30 mA. Grids are placed at 5cm from the silver target giving a coating rate of 0.4 nm/seconds requiring 20 minutes to reach the desired thickness of 500nm. The grid was then clipped in a standard Autogrid with a C-clip (Thermo Fisher Scientific Inc., US), and three reference marks were made on the Autogrid rim with a permanent pen: two dots aligned with the grid bars defining the future tilt axis and a third dot, on the side of the Autogrid, perpendicular to the tilt axis.

### Fine targeting in cryo-FIBSEM

The receiver silver grid was loaded at room temperature into the 45° pre-tilt shuttle under a binocular microscope, with the grid bars horizontally aligned such that the two reference dots were positioned horizontally (at 3 and 9 o’clock) and the third dot at the bottom (6 o’clock). This precise alignment facilitated subsequent serial sectioning of the lifted material.

The shuttle was then cooled down in the cryostation where the carrier was loaded and aligned using the engraved diamond marks (scratches) and gently cleaned with a fine brush and liquid nitrogen if ice contamination was present. After loading into the Aquilos 2 FIBSEM, an ion map was acquired following the procedure described in the Cryo-CLEM microscopy workflow section, and the fluorescent data was imported and aligned (Fig. 7H).

To avoid distortions from the elongated Z stacks and achieve precise targeting, the fluorescence Z stacks were reimported a second time, and the stage was navigated to each ROI. Ion beam snapshots were taken at each position, clearly visualizing the fiducial crosses, and the corresponding Z stacks were aligned using two alignment points avoiding distortion. Polar measurements (angle and distance) between fiducial crosses and features of interest were drawn in MAPS and these precise X-Y targeting coordinates are then drawn in xT software to place accurately the milling pattern to create the block (Fig. 7I-K).

### Axial-scaling factor to accurately target in Z

For accurate Z targeting, we accounted for the refractive index mismatch between the sample medium (water, n=1.33) and the immersion medium (N_2_ gas, n=1). Using established correction factors for the Leica Thunder microscope (NA=0.75, GFP emission λ=509 nm), we determined the actual depth of target (eg. 14µm) was 1.46 times greater than their apparent depth. An additional safety margin of 10 μm below the calculated target depth was implemented to protect the sample during undercutting.

### Block preparation

These preparations steps assume that the silver needle is already prepared following the Silver-Plated Needle Preparation, the EasyLift rod rotation check, and the silver needle milling steps. The ROI was centered on screen using MAPS software, the crosses are localized, and the precise coordinates of the block were defined using polar measurement from one cross. The target depth (eg. 14µm) was corrected using the 1.46 axial factor (Corrected target depth = TD = 14 x 1.46 = 20.44µm) plus the 10 μm safety margin. An opening trench was milled above the ROI, with the opening length (OL) calculated as: OL = (TD + 10) / tan (MA). The milling angle (MA), defined as the angle at which the undercut was performed beneath the block to release it from the carrier. For a corrected target depth of 20.44 μm, this resulted in an opening length of approximately 170 μm. The trench was milled using a 65 nA beam current. A secondary milling pattern was then applied at 50 nA to cut around the block, leaving a connecting bridge to the bulk material. The current was reduced to 15 nA when approaching the final block dimensions (50×80 μm) (Fig. 7J). The stage was then rotated to -70° and tilted to 17° (giving a 10° milling angle) to do the undercut of the block and the connecting bridge using a 5 nA rectangular pattern. The stage was then returned to the original position (110° rotation, 7° tilt) and the surface facing the opening trench was polished with a 5 nA cleaning cross-section pattern to make a smooth surface for needle attachment.

### Silver needle sample attachment

The silver-plated needle tip was flattened using a 5nA rectangular pattern prior to sample block attachment. The sample block surface was set at the coincident point, and the EasyLift needle was inserted and brought into contact with the block surface. Attachment was achieved through silver redeposition by placing a horizontal array of regular cross-sections (single pass, width 0.5 μm, lateral spacing 0.25 μm, height 2 μm, z-depth 4 μm) at 30 kV and 1 nA with the milling direction toward the silver needle. The block was released from the carrier by milling through the remaining bridge with a rectangular pattern (z-depth 10 μm, 30 kV, 5 nA) and lifted out of the carrier. The needle was retracted, the stage was lowered to remove the carrier from the field of view, the needle was reinserted, and the block width was reduced to 40μm, and the bottom was polished with a 5 nA current (Fig. 7L). Depending on the methods used (Serial Lif-Out, SOLIST) the sample is then respectively attached between grid bar (Fig. 10A) or deposited on the silver film of the receiving grid (Fig. 10B).

### SOLIST procedure

The SOLIST procedure was performed as described previously^9^. The first grid square for lamella deposition was positioned 6 squares to the left and 2-3 lines above the grid center. After setting the first grid square at coincidence point and 10° milling angle, the EasyLift needle was inserted, the block was lowered to slightly touch the silver film at the center of the grid square and a 42 μm wide, 600 nm high, and 3 μm deep rectangular milling pattern was used at 1nA to section a 5µm thick lamella from the block. After each cut, the block was raised, and the grid was moved by a relative displacement of -0.125 mm to the next grid square, repeating the process until the entire block was sectioned. This procedure allowed an 80 μm long block to yield approximately 13-14 lamellae.

### Lamella attachment on silver film

The grid was rotated 180° and tilted 7° perpendicular to the ion beam for attachment of all lamellae. Two arrays of regular cross section (single pass, width 2 μm, height 5 μm, z-depth 0.7 μm, 30kV, 1nA) were placed on the silver film on both sides of the lamella with the milling direction toward the silver film. Depending on the size of the lamella, one array can contain 4 to 8 milling patterns. The milling patterns were numbered following a milling sequence where the 1^st^ pattern is milled at the left of the lamella, the 2^nd^ at the right, the 3^rd^ at the left etc… to create a uniform, balanced and strong attachment of the lamella to the silver film.

### Lamella preparation to reduce curtaining

Once all the lamellae were attached, the gas injection system (GIS) was inserted perpendicular to the grid, and the flow was opened for 20 seconds to cover the top surface of each lamella. The front surface of the lamellae was then polished using cleaning cross-section patterns at 30 kV, 1nA (z-depth 4µm). The grid was then returned to the 10° milling position, and the GIS was reinserted for a 60 seconds deposition to cover the polished front surface of each lamella.

### Lamella thinning

Final thinning was performed using AutoTEM^TM^ Cryo software (Thermo Fisher Scientific Inc., US) starting at 1 nA to achieve an initial lamella thickness of 2 μm. Progressive thinning continued in steps: 1.4 μm thickness at 0.5 nA, 800 nm at 0.3 nA, 500 nm at 0.1 nA. Final polishing was performed manually at 50 pA using parallel rectangular milling patterns with an overtilt of 0.4-0.8° to thin down the back side of the lamella and achieve uniform thickness.

### Cryo-Electron Tomography

Data acquisition was performed on a Titan Krios G4 instrument operated at 300 kV equipped with a Selectris^TM^ X energy filter and a Falcon 4i camera (Thermo Fisher Scientific Inc., US). Tomographic data were collected using the Tomo5 software package (Tomo5 v5.2, Thermo Fisher Scientific Inc., US). Lamella-overview montages were acquired at a magnification of ×11,500 (pixel size, 2.236nm). Tilt series were recorded at a magnification of ×53,000, with a pixel size of 2.42 Å and stored in EER file format. Data collection followed a dose-symmetric tilt scheme with 2° angular increments, with 3 e⁻/Å² per tilt and a target defocus of −4.5 μm. The angular range spanned from −60° to 40°, resulting in a total accumulated dose of 150 e⁻/Å².

### Tomogram reconstruction

Frames were summed to get at least 0.8e-/px and motion corrected with alignframes command from IMOD package version 5.0.1^28^. Tomogram reconstruction was done with etomo from IMOD package version 5.0.1.

### Cold Trap Design and Implementation

A 1m long copper tube, 6mm outer diameter, 4mm inner diameter was annealed with a torch and quenched in tap water to soften it. It is then attached strongly in a vice with a 16mm cylindrical steel bar and manually rolled up around it to create a spiral (Fig. 8A). Two smaller brass tubes of 4mm outer diameter were brazed at the two ends of the copper spiral with silver solder. It is then cleaned with 10% sulfuric acid in water for 10min, then rinse several times with tap water and brushed with a brass brush to remove oxidation layer. It is then clean with deionized water and dried with acetone. The cold trap is then mounted in the heat exchanger of the cooling dewar by cutting and removing a small portion of the existing blue tube (blue tube length to remove = cold trap length – 2 small brass tube) and attach firmly to the main tube of the heat exchanger with two nylon cable ties as seen in Fig. 8B. During operation, the cold trap functions passively and doesn’t require any maintenance. The procedure below should be performed at the conclusion of extended microscope sessions, typically after 5 days of continuous operation to prevent trapped water from migrating into the microscope stage during system warm-up:

- Discontinue nitrogen gas flow.
- Extract heat exchanger from cooling dewar.
- Apply localized heating with a heat gun to the bottom tip of the cold trap to release the blue tubing connection.
- Restore nitrogen gas flow at 200 mg/s and heat the entire cold trap until complete moisture removal.
- Reestablish blue tubing connection to the cold trap.
- Reduce nitrogen gas flow to 10 mg/s for stage warming.

This maintenance protocol ensures safe system warm-up while preserving the microscope stage performance for subsequent sessions.

### Silver-Plated Needle Preparation

EasyLift^TM^ tungsten needles were silver plated using a RNG-1502 power supply (Bijoutil, Geneva, Switzerland) (Fig. 9A) which maintain a constant voltage/current and allow a continuous movement of the needle maintaining optimal plating conditions. The needle is first attached in an electric conductive holder and plunged in a degreasing bath containing Electrolytic degreasing type "A" (Bijoutil, Geneva, Switzerland) at a concentration of 70g/L in H2O, using two stainless steel anodes under a constant voltage of 2.5V and a current of 0.2A for 1min at RT. It is then rinsed several times in tap water following by rinsing in deionized water. The needle is then plunged in the silver cyanide solution "Argent W brillant" (Finishing, La-Chaux-de-Fonds, Switzerland) at a silver concentration of 25-40g/L using two fine silver anodes (Bijoutil, Geneva, Switzerland) under a constant voltage of 0.5V and a current of 0.1A for 10min at RT. The needle is then rinsed several times in tap water followed by rinsing in deionized water and finally dried at room temperature. A thickness measurement check was done under a binocular using a 300 square mesh copper grid (space between the grid bar is around 50-60 µm) attached with a transparent tape on a glass slide. The needle tip was aligned onto the grid to estimate the thickness by comparing it with the width of a grid square.

### EasyLift^TM^ rod rotation check

The 90° rotation check of the EasyLift^TM^ rod was done at room temperature with the microscope chamber open. In the starting position, the red mark of the EasyLift^TM^ connector should be upward (Fig. 9B <u>1</u>) and the cooling braid clamp should be on the right (Fig. 9B 2). Then, the EasyLift^TM^ rod was slowly rotated clockwise (CW) and continuously checked to see if the copper cooling braid had enough freedom to reach 90° of rotation (red mark of the EasyLift^TM^ connector to the right, Fig. 9B 3) without tension on the cooling braid and its clamp (Fig. 9B 4). Then the rod was rotated back counterclockwise (CCW) slowly to 0° (red mark upward Fig. 9B 1) while checking the movement of the cooling braid. This rotation should not make too much tension on the cooling braid, and the clamp should stay in place while rotating.

Once this rotation check was done, the cooling braid clamp was open, the EasyLift^TM^ rod was removed from the microscope to install the silver needle and finally reinstalled in the microscope. The rotation of 90° CW/CCW was then done under high vacuum and cryo-condition for leak checking (the vacuum value of the chamber can be checked when rotating 90° CW/CCW).

### Silver needle milling steps

The preparation can be performed at room temperature or under cryogenic conditions. The needle is inserted and moved to the coincidence point between the electron beam (EB) and the ion beam (IB) (Fig. 9C 1-2). A polygonal pattern is drawn at the needle tip to remove the top half of the needle material over a length of 250 µm (Z-depth 40 µm, 30 kV, 65 nA) (Fig. 9C 3-4). Using the EB view the silver layer thickness around the tungsten core was measured and should be around 25-30 µm (Fig. 9C 5). A rectangular tilted pattern is drawn below the needle to creates a flat bottom face over a length of 250 µm (height 1 µm, Z-depth 40 µm, 30 kV, 15 nA) (Fig. 9C 6). While scanning with the IB (1536x1024, 200 ns, 10 pA) the EasyLift^TM^ rod is rotated 90° CW until the red mark of the EasyLift^TM^ connector is positioned to the right, exposing the bottom face of the needle in the IB view. Two polygonal patterns are drawn to thin down the needle to 25 μm (Z-depth 40 µm, 30 kV, 65 nA) (Fig. 9C 7-8). The EasyLift^TM^ rod was then rotated 90° CCW to the original position by placing the red mark of the EasyLift^TM^ connector to the top and a polygonal pattern was drawn to remove the remaining tungsten and achieve 17 μm thickness (Z-depth 40 µm, 30 kV, 15 nA) (Fig. 9C 9). The EasyLift^TM^ rod was rotated 90° CW and two rectangular tilted patterns were drawn to thin down the needle to 17 μm in the second dimension (Z-depth 40 µm, 30 kV, 15 nA) (Fig. 9C 10). The EasyLift^TM^ rod was finally rotated 90° CCW to its original position.

The final needle had a length of 250 µm and 17 μm thickness in both dimensions, given a horizontal width of 25 μm for attachment to the sample (9C 11-12)

## Conclusion

Our study demonstrates that recent advancements in cryo-ET significantly enhance the structural analysis of plant root tissues while maintaining consistency with traditional chemical fixation methods. The ability to preserve native cellular structures with minimal artifacts offers new insights into plant cell architecture and molecular organization. Importantly, our findings confirm that cryo-ET does not invalidate prior models but rather refines them, allowing for a more detailed examination of cellular processes such as suberin deposition and Casparian strip formation.

By comparing full cryo-workflows with resin embedded chemical fixation, we highlight the importance of selecting the most suitable technique based on specific research questions. Cryo-ET excels in preserving osmotic balance and ultrastructural integrity, enabling high-resolution investigations of plant cell wall dynamics. However, traditional fixation remains valuable for large-scale and quantitative analyses.

As cryo-ET methodologies continue to evolve, they will provide unprecedented opportunities to explore plant cell biology at the molecular level. Future studies integrating correlative light and electron microscopy approaches will further bridge the gap between structural and functional analyses, advancing our understanding of plant development and adaptation.

## Supplementary material

All the movies are accessible through this link: https://filetransfer.dcsr.unil.ch/message/Bh2Tf0LuykFPYOQtAB7P2N

## Acknowledgements

This work is based on the publications of two Nature Methods papers that have inspired and guided us during this work. We thank the authors of these key papers for the detailed protocols that allow us to implement the technique on the EM facility and motivate us to develop them. We thank Alexander Myasnikov and Bertrand Beckert from the Dubochet Center of Imaging, UNIL and EPFL, Lausanne, for their training and supervision for cryo-ET acquisitions. We thank Christian Zimmerli for fruitful discussions during the development of the method. We thank Christopher Thompson (Thermo Fisher Scientific) for the discussion about metals sputter rates that led us to the idea of making the silver needle. We thank the company Delmic that has loaned the Ceres clean station that was instrumental to reduce the ice contamination during our transfers.

